# N6-methyldeoxyadenosine is a transgenerational epigenetic mark protecting DNA from deletion

**DOI:** 10.64898/2026.02.13.705781

**Authors:** Xiaodong Li, Sarah Allen, Liping Lyu, Sebastian Bechara, Cristina Engeroff, Alan Hendrick, Mariusz Nowacki

## Abstract

DNA N6-adenine methylation (6mA) is emerging as an important DNA modification present in all eukaryotic kingdoms, where its roles mostly involve transcriptional regulation and chromatin modification. Here, we demonstrate a distinct role for 6mA in *Paramecium tetraurelia*. We find that it is a trans-generational epigenetic mark which distinguishes host DNA from invasive elements, protecting the former. During development, Paramecium deletes around 25% of its germline genome, removing TEs, repetitive sequences and transposon-derived elements, to form a functional somatic genome. We show that 6mA is present in the Paramecium germline nucleus, where it is preferentially located on host DNA and excluded from TEs. Ectopic addition of 6mA to germline DNA causes failure to excise TE-derived sequences, leading to lethality. 6mA presence on a synthetic TE-derived element is sufficient to block its excision during development. These effects are inherited via sexual reproduction, demonstrating their trans-generational capacity.

## INTRODUCTION

Trans-generational epigenetic inheritance is the transmission of genetic information from parent to multiple generations of offspring without altering the DNA sequence itself. In sexually reproducing organisms, trans-generational epigenetic marks involve modifications to the germline, as this is the origin of the F1 and all future generations of offspring. To date, the known trans-generational epigenetic marks include cytosine CpG methylation (plants, protozoa, mammals)^1,2,3,4,5,6^ histone modifications (plants, protozoa, invertebrates, mammals)^7,8,9,10^, prions (fungi)^11^ and small RNAs (plants, protozoa, invertebrates)^12,13,14^. The idea that changes to our genomes acquired during our lifetimes could be passed on to our offspring is fascinating and the subject of much debate, as it is often difficult to separate genetic effects on the epigenome from true epigenetic inheritance^15^.

Ciliates possess a number of features that make them an excellent model system for trans-generational epigenetic inheritance. Firstly, they exhibit nuclear dimorphism, meaning that germline and soma are separated into two distinct nuclei within the same cell^13,16^. The germline nucleus is termed the micronucleus, or MIC, due to its relatively small size. It exists in a transcriptionally silenced state during normal vegetative growth^17^. The somatic nucleus is present at high copy number, making it larger in size, and is hence termed the macronucleus (MAC). It is transcriptionally active and acts as the working nucleus for vegetative growth. Vegetative growth refers to the non-sexual part of the ciliate life cycle, during which ciliates feed and divide. The MIC divides by mitosis at each cell division, while the MAC divides by amitosis. During sexual development, the micronucleus undergoes meiosis, and two of the resulting haploid pronuclei fuse to form the zygotic nucleus, which then divides twice to form four diploid nuclei. Two of these will form new micronuclei to continue the germline, and the other two will form the macronuclei for two daughter cells. In conjugation, which is the sexual process for most ciliates, one of the haploid pronuclei is exchanged between the maternal cell and another cell of a different mating type, prior to fusion and formation of the zygotic nucleus. As the new macronuclei are formed, the old macronucleus in the parent cell (the maternal macronucleus) is degraded and eventually disappears. The process of forming a new macronucleus from a micronucleus is called development, and is a remarkable event in which the genome is amplified up hundreds of times (around 800-1000 times in *Paramecium tetraurelia*)^18^. Meanwhile transposons, repetitive DNA and transposon-derived sequences termed Internal Eliminated Sequences (IESs) are removed from the genome. Thus, the macronuclear genome is shorter than the micronuclear genome, but is more streamlined, having removed all DNA sequences deemed threatening or unnecessary for general life. To distinguish between genomic DNA required in the new macronucleus (MAC-Destined Sequence or MDS) and unwanted interrupting DNA (IES, transposon etc.), ciliates employ a range of epigenetic mechanisms. These mechanisms involve information transfer from the maternal macronucleus to the developing daughter macronucleus, and include small RNA pathways, RNA-guided histone modifications and in some species, long guide RNAs^8,19–24^. Thus, ciliates pass epigenetic information from maternal soma to daughter soma in a paradigm of epigenetic trans-generational inheritance.

In *Paramecium tetraurelia*, there are around 45,000 IESs scattered throughout the genome, the vast majority within protein-coding genes or regulatory sequences^25,26^. Therefore their excision must be highly precise, as any inaccuracy would lead to indel mutations, potentially disrupting the reading frame of the genes in which they are located. IESs are excised by a domesticated PiggyBac-like transposase named PiggyMac, which cleaves at an obligate TA dinucleotide at each end of an IES, removing the IES and leaving one TA behind^27^. Apart from these flanking TA dinucleotides, *Paramecium* IESs have a weak consensus of TAYAG inverted repeats at each end^25,26,28^. This is, however, not enough information for the PiggyMac to correctly and accurately recognise all 45,000 IESs in the genome. So how are IESs recognised? One mechanism is the scanRNA pathway, in which small RNAs termed scanRNAs are generated from the MIC and compared against the maternal MAC. Those scanRNAs that fail to find matches in the maternal MAC are sent to the developing MAC to target MIC-limited sequences for removal. This is a very elegant and well-studied model, however it only accounts for the recognition of around 15% of IESs. The remainder are seemingly recognised without any need for scanRNAs, and are termed non-maternally controlled IESs. The current model for non-maternally controlled IES excision posits that their recognition depends on a variety of factors, none of which are alone sufficient to identify IESs but which, when taken together, can provide the cell with enough information to reliably identify IESs and target them for excision. These factors include the flanking inverted repeats including the obligate TA, the distance between these flanking TAs^26^, nucleosome positioning in the developing MAC^29^, and a recently-discovered protein that aids in the recognition of a subset of IESs^30^. However, even taken together, these factors do not satisfactorily explain how a *Paramecium* cell can identify 45,000 IESs within a 100 megabase genome, over a period of a few hours, with accuracy down to the single base pair. In addition, non-IES sequences are reliably retained, with erroneous (or’cryptic’) excision of MAC genomic sequences occurring at vanishingly low levels, despite the fact that the genome is AT-rich (around 72%)^31^ and TA dinucleotides are therefore highly abundant.

DNA N^6^-adenine methylation (6mA) is a DNA modification that has gained attention in recent years, as technological advances allowed its detection in model organisms such as *C. elegans*, *Drosophila*, zebrafish, plants, and even mice and humans^32–36^. It is the primary DNA modification in ciliates^37^, where it has been shown to have a positive association with transcription in *Tetrahymena*^38,39^, and a negative association with histone occupancy in both *Tetrahymena* and *Oxytricha*^40,41^. In worms and flies, 6mA is also associated with gene regulation and chromatin modifications, particularly during development^42,43^. In vertebrates, 6mA is a relatively rare modification and its functions are not yet well characterised, though 6mA levels appear to fluctuate with development and various stressors^44^. So far, 6mA has not been well-characterised in *Paramecium tetraurelia*, and has not been shown to have a role in trans-generational epigenetic inheritance. We wished to discover whether 6mA may act as a marker helping Paramecium to distinguish MDS from IES during development.

## RESULTS

### 6mA preferentially located on non-IES sequence in Paramecium germline

Initially we wished to establish what DNA modifications are present in *Paramecium* and whether any of them are differentially present in MDS versus IES. Since PiggyMac (PGM) is the enzyme that excises IESs, knockdown of PGM (PGM-KD) leads to the retention of IESs in the MAC following development, leading to a non-functional MAC and cell death. We extracted MAC DNA from wild-type and PGM-KD cells and performed mass spectrometry analysis to quantify all DNA modifications known in biology. No DNA modifications were detected at significant levels except 6mA, which was present in approximately 0.6% of all adenines (see Figure 1A). In PGM-KD DNA, the relative abundance of 6mA was approximately 0.75x of WT DNA. This is consistent with DNA preferentially being methylated on non-IES sequences, or MAC-Destined Sequences (MDS), as these are relatively less abundant in PGM-KD DNA where IESs are retained. 5-methylcytosine (5mC) was measured at just above the detection limit of the mass spectrometer, consistent with previous work that has shown 5mC to be absent in Paramecium^45^. The trace levels detected in our experiment are likely to come from low levels of feeding bacteria present in the media, or endosymbiotic bacteria within the Paramecia.

**Figure 1.**
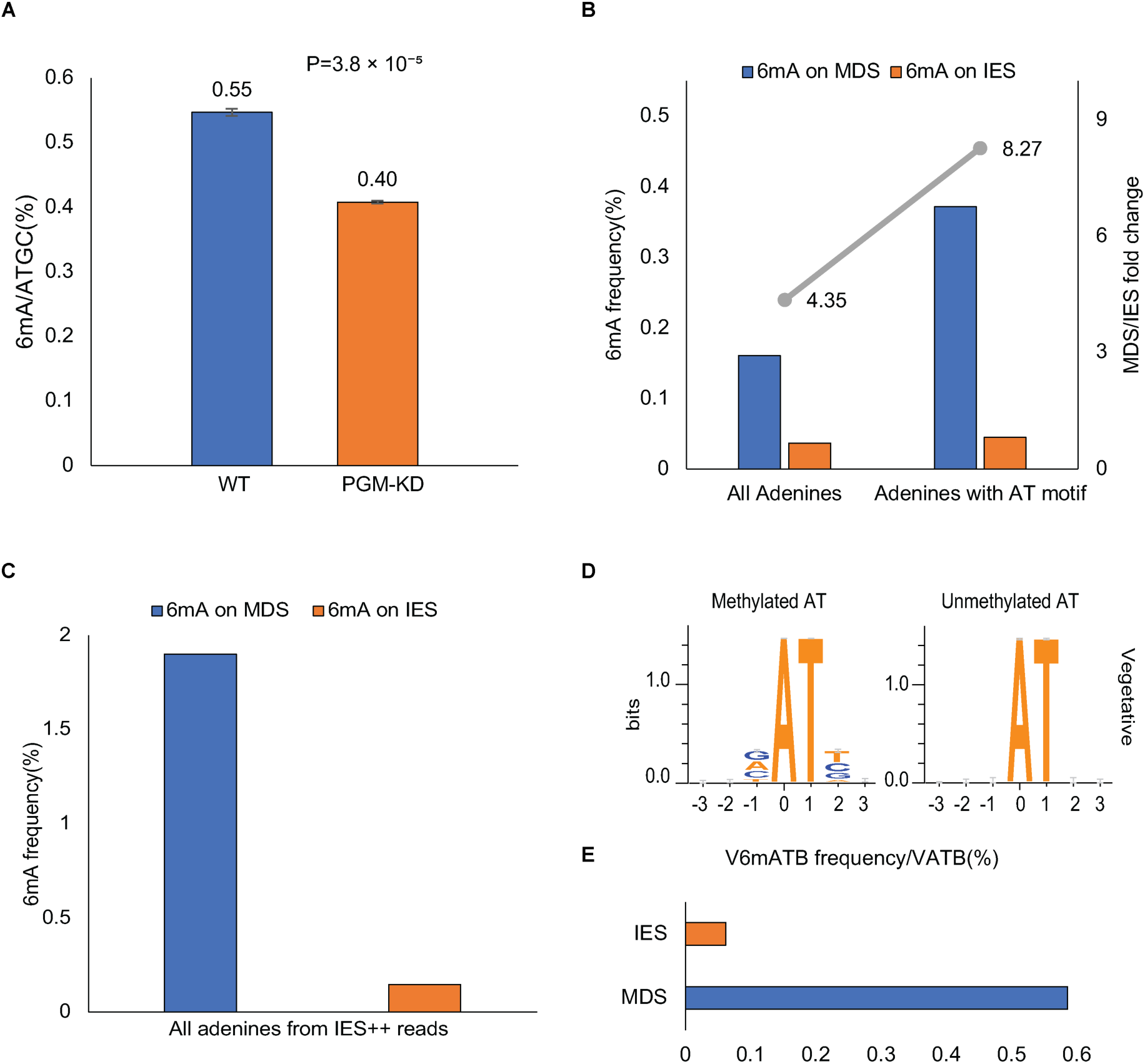
Mass Spectrometry and SMRT sequencing results showing 6mA frequencies on MDS versus IES. (A) Values are shown as a percentage of total Adenines. 6mA levels were significantly reduced in PGM-KD compared with WT (0.45 ± 0.02 vs 0.61 ± 0.01, mean ± SD; Welch’s two-sample t-test, p = 0.00086; n = 3) (B) vegetative MIC DNA reads were selected by including only reads containing at least two IESs. 6mA frequencies are shown for all adenines, and for adenines within an AT motif. Fold change between MDS versus IES is indicated by the grey bar and the Y axis on the right. (C) The same analysis on DNA from developing MACs in PGM-KD cells. This DNA should contain the 6mA marks carried through from the MIC, but vegetative transcriptional 6mA marks could also be beginning to be established. (D) Sequence logos of AT dinucleotides either methylated or unmethylated, from the MIC DNA in A. (E) Analysis of 6mA status of VATB tetranucleotides in MIC DNA, on IES versus MDS.

Having established that 6mA is present in higher levels in WT DNA than PGM-KD DNA, we wished to know whether this was due to preferential methylation of MDSs as opposed to IESs, or some other reason, such as gene misregulation in the compromised PGM-KD MAC. We performed SMRT PacBio sequencing on WT vegetative cells, then selected reads that contained at least two IESs for analysis. We reasoned that in vegetative WT *Paramecium*, any genomic DNA containing at least two IESs must come from the MIC, since the MAC will be devoid of IESs. Since the MAC is present at ∼800n, compared to 4n for the two MICs, the number of reads obtained by this method was small. Nonetheless, as shown in Figure 1B, 6mA was enriched 4-fold on MDSs compared to IESs. We observed that the majority of 6mA was located on AT dinucleotides (84%). If the methylated adenine in an AT dinucleotide is an epigenetic mark distinguishing MDS from IES, we would expect to see a stronger bias towards MDS methylation when exclusively comparing methylated adenines in AT dinucleotides rather than all adenines. This is indeed the case, as shown in Figure 1B. When comparing 6mA exclusively located in AT dinucleotides, the bias towards MDS rather than IES increases to 8-fold. This supports the notion of 6mAT being an epigenetic marker for MDS.

In the *Paramecium* genome annotation we used for the analysis, a small proportion of IESs are mis-annotated, probably due to differences between IESs in different strains of *Paramecium*^46^. These “non-IESs” are easily recognisable, being annotated as 100% retained in WT cells, or as 100% excised in PGM-KD (see Supplementary Figure S1F and G). Since there are relatively few of them we usually ignore them, but since the number of reads in our initial analysis was low, we decided to remove them from the analysis to see whether it had an effect on the results. Analysis of 6mA in IES versus MDS after removal of mis-annotated IESs is shown in Supplementary Figure S1A. The effect is similar to the results of the global analysis, but stronger, confirming that 6mA placement negatively correlates with excision. We decided then to repeat the analysis including reads containing only one IES. These are likely to come from the MIC but with a lower certainty, as occasional random retention of single IESs could occur in the MAC as well. When including single IES+ reads, the results of the analysis look similar but with a stronger preference for 6mA on MDSs (Supplementary Figure S1B-S1E). When further removing the mis-annotated IESs from these reads, the effect was stronger still, with a 24-fold over-representation of 6mA on MDS compared to IES.

One notable observation from our data is that the proportion of 6mA in MIC DNA is low compared to the result of the mass spectrometry. The reason for this is probably twofold: the mass spec was performed on post-autogamous cells and therefore contained a high proportion of MAC DNA, which may have much higher levels of 6mA than MIC DNA. Secondly, in our analysis of the SMRT PacBio sequencing, we used the highest stringency threshold for calling modifications, as we wished to be as certain as possible that any modifications identified were true modifications and not false positives. Therefore, it is likely that a number of 6mA modifications were missed due to low certainty in the analysis. The true proportion of 6mA in the MIC is probably somewhat higher than that shown in figure 1B (0.1-0.2%), but lower than the mass spec levels (0.6%) which contain transcriptional 6mA modifications.

We were curious to see whether there is a preference for the nucleotides surrounding methylated ATs. We generated sequence logos for AT dinucleotides in which the A is methylated, compared to all AT dinucleotides. As shown in Figure 1D, the nucleotide preceding a methylated AT shows a preference for non-T, and the following nucleotide has a preference for non-A. When unmethylated ATs are included in the analysis, this preference disappears. Therefore, TA dinucleotides on both strands are excluded from 6mA methylation. This is consistent with 6mA having a role in distinguishing IES from MDS, as TA dinucleotides flank every IES, and TAYA is part of the preferred consensus internal repeat on IESs. If 6mA is an epigenetic marker identifying MDSs for retention in the developing MAC, then IES ends, where cleavage takes place, should be strongly avoided as methylation targets.

To ensure that the preferential methylation of MDSs is not simply a result of them having fewer TA dinucleotides, we repeated the analysis in Figure 1B, but only included VATB tetranucleotides. As shown in Figure 1E, the preference for 6mA in MDS over IES remains, and is indeed stronger than in all ATs, with 0.6% of VATB tetranucleotides being methylated on the A in MDS, compared to only 0.02% in IES. Whether this sequence preference is due to the methyltransferase or for another reason is as yet unclear. Intriguingly, the same sequence preference is seen in *Oxytricha* MAC 6mA^40^ and *Chlamydomonas reinhardtii*^47^, but not in the more closely related ciliate *Tetrahymena*^38,39^.

Since the number of MIC reads obtained from total DNA is relatively low due to the difference in ploidy between the different nuclei, we wished to repeat our analysis on a larger dataset. In PGM-KD, cells are unable to excise IESs during sexual reproduction, and the new MAC copy number is amplified with IESs still present. Therefore, the new MAC in PGM-KD cells post development can be considered a representation of the MIC in terms of sequences present. A feature of *Paramecium* is its ability to go through autogamy, which is sexual reproduction in the absence of a partner. Instead of swapping a haploid meiotic pronucleus with another cell, *Paramecium* can simply fuse two haploid pronuclei within the same cell, and then continue with development as if conjugation had occurred^13^. We allowed PGM-KD cells to go through autogamy and then isolated macronuclei in the late stages of development. We then performed PacBio SMRT sequencing on the DNA obtained. For analysis we once again chose sequencing reads containing at least two IESs on the same read, to ascertain that what we were looking at was newly formed MAC and not the maternal MAC. The maternal MAC is still in the process of being degraded and so its DNA is still present in large amounts in the cell, but it does not contain IESs as it was formed one generation earlier, in the presence of PGM. Results are shown in Figure 1C. Similarly to in vegetative MIC DNA, 6mA was preferentially located on MDSs, approximately 12-fold more than on IESs. We acknowledge that 6mA may also be a transcriptional marker in the vegetative MAC, similarly to in *Tetrahymena*^38^. Therefore, it is not impossible that vegetative methylation marks in the MAC are already being established in late development, obscuring any marks carried from the MIC to the new MAC. Nonetheless, the pattern of methylation looks similar between vegetative MIC and PGM-KD developing MAC.

### Aberrant methylation in the germline nucleus leads to retention of IESs in progeny

To investigate the function of methylation in *Paramecium*, we decided to investigate the effect of ectopic methylation of MIC DNA to the cell. We targeted a non-specific 6mA methyltransferase, Hia5^48^, to the vegetative *Paramecium* MIC by fusing it to the MIC-specific centromeric histone H3 (CenH3) protein^49^, and microinjecting into the *Paramecium* MAC. Microinjection into the MAC is the standard method for introducing a transgene into *Paramecium*, as *Paramecium* will retain the introduced gene in its MAC by telomerising it and treating it like a microchromosome, expressing it and replicating it with each division, until the next sexual reproduction, whereupon the gene is lost^50^. *Paramecium* MIC chromosomes are holocentromeric, so our protein fusion should be distributed along the chromosomes where it can methylate DNA indiscriminately. Initially, we fused Hia5 and GFP to the CenH3 C-terminus, to confirm visually by GFP fluorescence that the construct is targeted to the vegetative MIC. As shown in Figure 2A, after microinjection of the construct, MICs can clearly be seen to contain GFP, while MACs do not.

**Figure 2.**
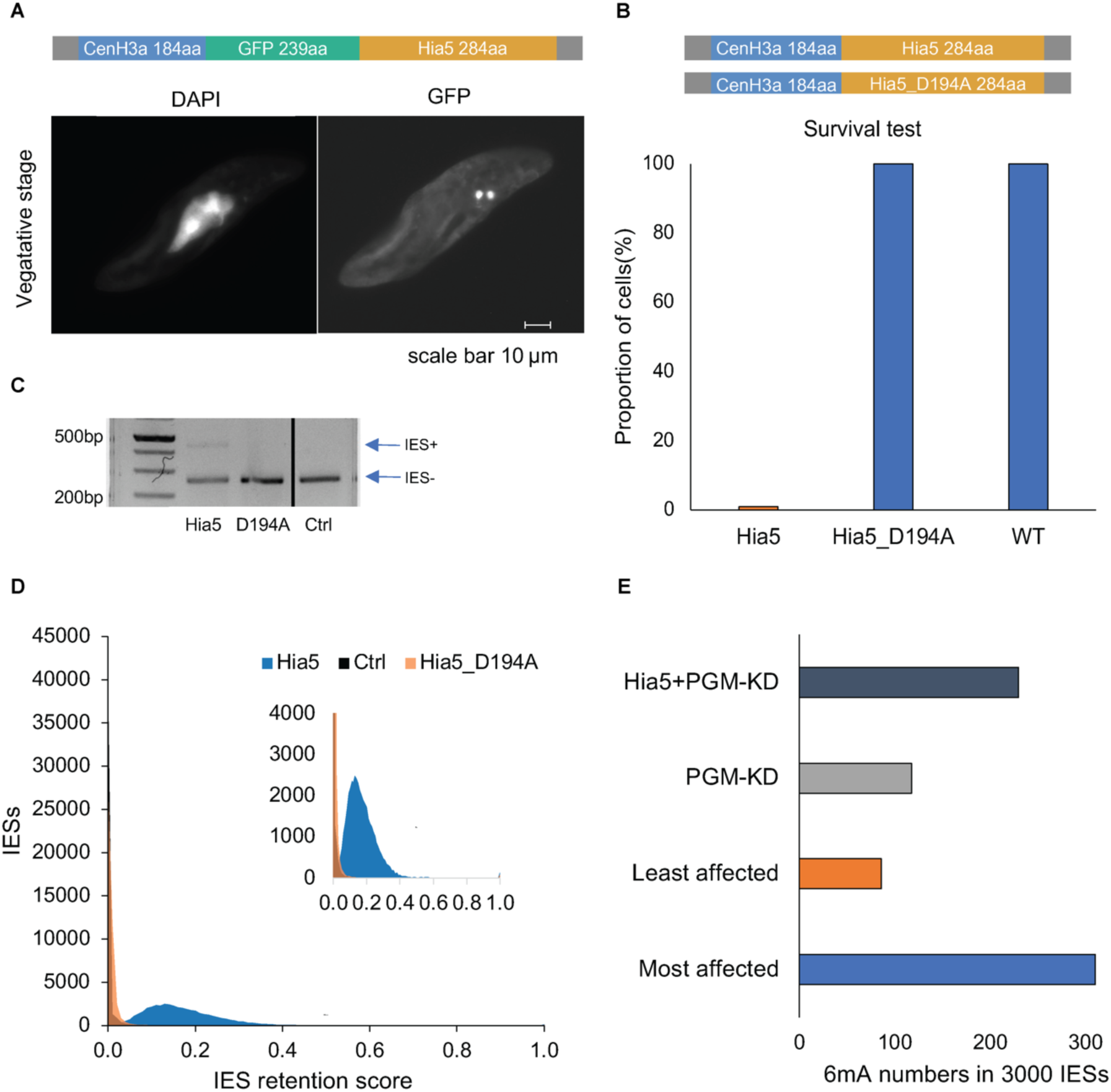
Ectopic Hia5 expression in the vegetative MICs causes lethality and IES retention after sexual reproduction. (A) Schematic of the CenH3-GFP-Hia5 construct and fluorescence microscope image showing GFP localisation of the fusion protein in vegetative cells. DAPI staining shows the localisation of the vegetative MAC and MICs. (B) Schematic of the CenH3-Hia5 construct without GFP, and of the catalytic mutant D194A. Survival test following autogamy shows lethality of the catalytically active Hia5 but not of the catalytic mutant, with WT cells as control. (C) IES retention PCR of DNA extracted from post-autogamous cells. (D) Global IES retention post-autogamy in Hia5-expressing cells, WT cells and Hia5-D194A-expressing cells. IES retention score is the proportion of reads with retention relative to the total number of reads for a given IES^46^. (E) 6mA marks/3000 IESs. 6mA marks on the 3000 IESs with the highest retention scores in the Hia5-expressing cells (“most affected”) were counted and compared to the number of 6mA marks on the 3000

IESs with the lowest retention scores in Hia5-expressing cells (“least affected”). For comparison, average 6mA marks per 3000 IESs are shown for PGM-KD and in Hia5+PGM-KD.

Having confirmed visually that the construct is successfully targeted to the MICs, we removed the GFP gene in case it interfered with the Hia5 function. Additionally, we generated a Hia5-CenH3 construct containing a mutation in which the catalytic aspartic acid residue D194 is mutated to an alanine (Hia5_D194A). This abolishes the methyltransferase function of the enzyme^48^. We then microinjected the Hia5-CenH3 constructs into separate Paramecium cells and allowed them to grow vegetatively for ∼16 divisions before starving to initiate autogamy. Following autogamy, individual cells from the F1 were picked into separate wells and fed to assess growth and survival. As shown in Figure 2B, lethality was 100% in post-autogamous cells from the Hia5-injected parental cells. In WT and Hia5_D194A-injected cells, lethality was 0%. If 6mA is an epigenetic mark that protects MDS DNA from excision, we hypothesised that aberrant methylation from Hia5 may be preventing IESs from being excised, leading to IES retention and therefore disruption of genes, and cell death. To test this, we performed PCR with primers spanning known IES regions. IES retention causes the appearance of a longer (IES+) band in addition to the shorter band that is present in WT cells. As shown in Figure 2C, an IES+ band is visible in the progeny of the Hia5-injected cells, but not in the progeny of the WT or Hia5_D194A cells. This confirms that 6mA added by the active Hia5 enzyme, but not the mutant version, leads to retention of IESs in F1, consistent with the idea that 6mA is a marker for retention in the new MAC. To further investigate IES retention in the new MAC of progeny from Hia5-injected cells, we performed deep sequencing of MAC DNA from Hia5, WT and Hia5_D194A progeny. Only Hia5 progeny showed IES retention (Figure 2D). IES retention is measured by the IES Retention Score (IRS) which simply calculates the number of IES+ reads divided by IES-reads. As shown in Figure 2D, IESs are retained globally between roughly 5% and 40%, as compared to WT and Hia5_D194A cells, which show no IES retention.

We wished to assess whether the ectopic methylation of IESs in the presence of Hia5 is associated with their retention. To do this, we needed an unbiased representation of the methylation status of all IESs following Hia5 expression, compared to WT. In Hia5-expressing cells, the majority of IESs were still excised to a certain degree, and their methylation status could thus not be determined. Therefore, we performed PGM-KD on Hia5-expressing cells (Hia5+PGM-KD). PGM-KD, as described, leads to global retention of IESs regardless of their methylation status. Performing SMRT PacBio sequencing on DNA from these cells provides us with information about which IESs are methylated (and which are not) in the Hia5-expressing MIC, regardless of their excision. To look at whether IES methylation correlates with retention, we chose the 3000 IESs with the highest retention scores in the Hia5 progeny, and the 3000 IESs with the lowest retention scores. We then assessed their average methylation levels in Hia5+PGM-KD DNA. Stronger IES retention after Hia5 expression was associated with higher 6mA levels, although levels remained low overall. Notably, the 3000 least-affected IESs had an average methylation level similar to the baseline PGM-KD-only, implying that they escaped methylation by Hia5.

To ascertain that the Hia5 phenotype is not due to loss of small RNAs due to for example blocked transcription in the meiotic MIC, we performed small RNA sequencing on WT and Hia5-expressing autogamous cells. As shown in Supplemental Figure S2B, scanRNAs were not affected. We also wished to determine whether the IES retention phenotype could be due perturbation of genes involved in the genome rearrangement pathway, rather than directly due to methylation of IESs. We performed correlation analysis to see whether the IES retention was similar to in knockdown of any known genes. As shown in Supplementary Figure S2A, IES retention following Hia5 expression does not look similar to any known genes involved in genome rearrangement.

### 6mA efficiently blocks excision of an IES in vivo

Our next aim was to ascertain whether 6mA marks can directly block excision of an IES in *Paramecium*. To do this, we designed synthetic constructs containing an IES flanked by two MAC sequences from elsewhere in the genome. We also included the endogenous 10 bp on either side of the IES to ensure that any regulatory sequences were present. The goal was to create an IES that would be recognised and excised by the cellular machinery, but which could also be identified and PCR-amplified independently of the endogenous IES. We then had two versions of the construct synthesised, one with added bidirectional 6mA marks at four positions along the IES, and one without 6mA. The two versions of the construct were also differentiated by a 3bp’barcode’ tag to later distinguish between them by sequencing (see Figure 3A).

**Figure 3.**
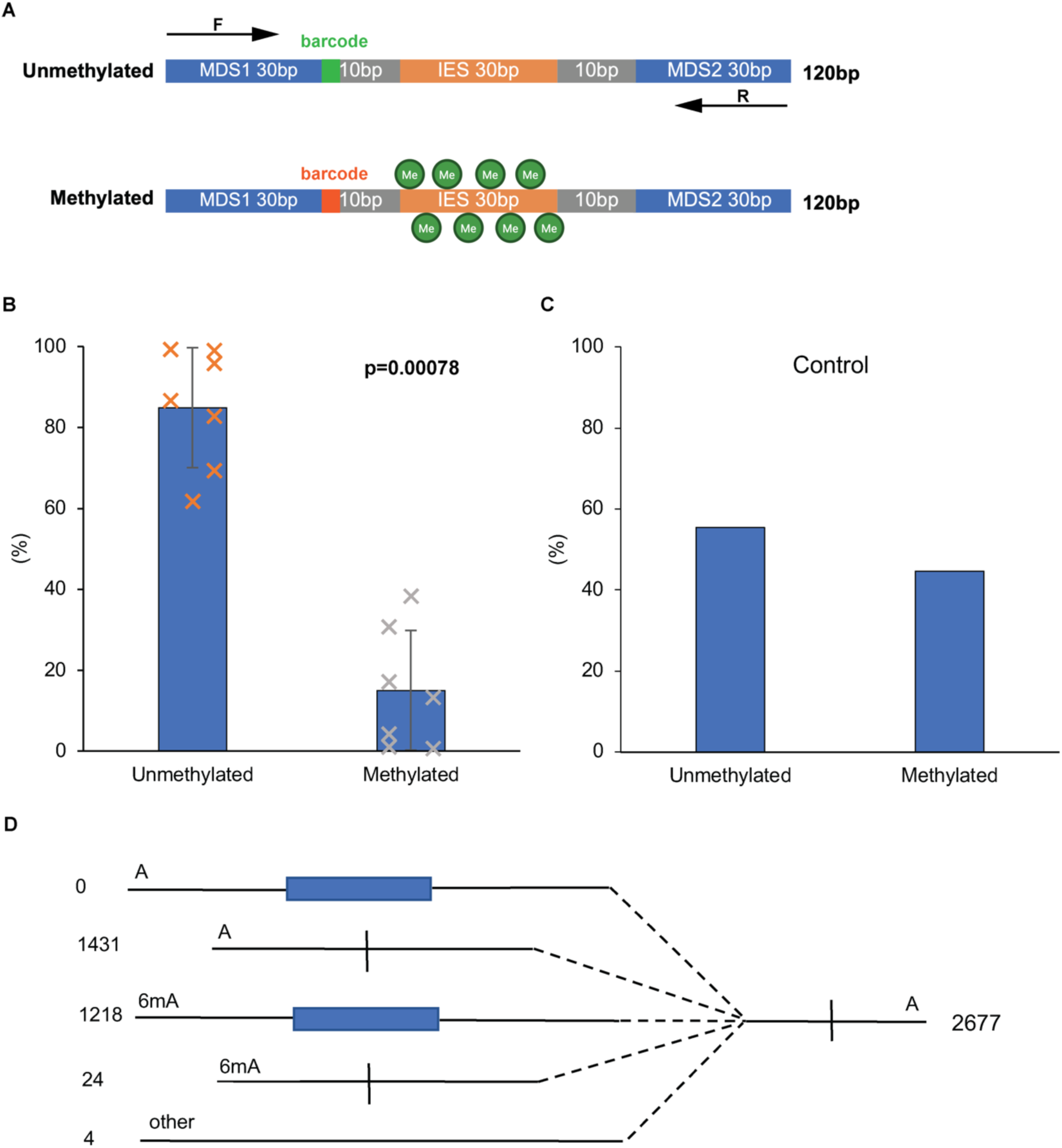
Injection of methylated and unmethylated IESs to compare their excision. (A) schematic of the two injected constructs. The MDSs and IESs are both endogenous Paramecium sequences but from different genomic locations, to allow for easy amplification of injected constructs by PCR. The IESs are flanked by 10 bp on each side of their endogenous flanking sequence (grey in the schematic). The barcodes allow for identification of the methylated versus unmethylated versions by sequencing. (B) Reads on which the IES was excised were sorted by methylation status. The graph shows % unmethylated versus % methylated. A paired two-tailed t-test revealed a highly significant reduction in excision in the methylated group compared to the unmethylated group (Unmethylated: 84.9% ± SD, Methylated: 15.1% ± SD; t = 6.25, df = 6, p = 0.00078). (C) PCR and sequencing of the input DNA as in B, to check the proportions of methylated versus unmethylated DNA in the injections. (D) analysis of concatenated DNA segments to assess whether methylated DNA is imported into the MAC as efficiently as unmethylated.’A’ denotes unmethylated construct,’6mA’ denotes methylated. The right hand side is the total reads for which an unmethylated excised construct is concatenated at its 5’ end to another piece DNA. The left hand side shows the proportions of different sequences to which it was concatenated. Methylated DNA construct was present and excised at the same level as in the total DNA analysed (B), indicating that methylated DNA is imported as efficiently into the MAC as unmethylated DNA.

We injected a mixture with equal concentrations of each version of the construct into Dcl5-silenced autogamous Paramecium cells. Short (<∼200 bp) DNA sequences injected during mid to late autogamy have been shown to enter the developing MAC during the ongoing genome rearrangement process^51^. Dcl5 is a gene involved in the production of iesRNAs, which are a secondary class of small RNAs that are produced from concatenated excised IESs during the later stages of sexual development. The reason we silenced this pathway was because injected DNA can enter the iesRNA pathway which creates a positive feedback loop, ensuring excision of all copies of an IES. Previously we have shown this positive feedback loop to be capable of triggering excision of non-IES sequences when large quantities of DNA are injected^51^. We wished to avoid the possible confounding influence of the iesRNA pathway feedback loop and focus on whether 6mA marks can prevent excision under normal physiological circumstances.

Following injection, we allowed the cells to complete around 10 divisions before harvesting their DNA and performing PCR to amplify the synthetic construct. We sent PCR products for deep sequencing to analyse what proportion of a) methylated and b) unmethylated IESs were successfully excised. We identified the methylation status of the injected construct based on the barcode tag previously mentioned. Results are shown in Figure 3B. Seven individual injected cells were grown up and their DNA sequenced. The average excision level for the unmethylated IES was around 85%, and for the methylated IES it was around 15%, and a paired two-tailed T-test confirmed a highly significant reduction in excision on the methylated construct (p=0.00078). There is considerable variability between the seven different samples, with some displaying close to 0% excision on the methylated version and one displaying over 30% excision. We suspect that this variability is due to variability in the timing of the injection and the efficiency of the Dcl5 silencing, which varies from cell to cell and also decreases over time. If excised IESs from the injected DNA are concatenated into iesRNA templates, the iesRNA pathway would generate vast numbers of iesRNAs targeting that particular IES, as the injected DNA is present at high levels. This could be a strong enough signal to override the methylation marks. If cells were injected later during development, as the Dcl5 silencing effect was wearing off, then the injected DNA could enter the iesRNA pathway. Nonetheless, the majority of injected cells display much lower excision rates on the methylated than the unmethylated constructs. We also performed PCR and sequencing on the input mixed DNA to ascertain that the two versions of our construct were injected in equal proportions. This was not in fact the case (Figure 3C), as the unmethylated version was present at 55.4% in the input DNA, versus 44.6% for the methylated. These levels are, however, comparable - and should have no bearing on the relative efficiency of excision of the two versions.

We then wished to ascertain that the difference in excision efficiency was due to a preference by the excision machinery, and not because the methylated construct entered the nucleus at lower efficiency than the unmethylated, and was therefore not exposed to the excision machinery at all. To address this problem, we made use of the fact that Paramecium tends to concatenate injected DNA into longer strings, each one containing multiple copies of an injected sequence^52^. This concatenation occurs at random so, in our case, we could expect methylated and unmethylated constructs to be concatenated together with high frequency. We selected reads from one of the injected cells that had had the greatest difference in excision efficiency between methylated and unmethylated injected IESs, as we reasoned that if there is a difference in import efficiency then it would be most readily detectable there. We selected all reads that had an unmethylated construct that had been successfully excised, and to which another sequence was concatenated on its 5’ end. We reasoned that these reads had surely entered the developing MAC, as otherwise the IES on the 3’ unmethylated sequence would not have been excised. The total number of such reads was 2677. We then examined the 5’ connected reads to determine what they were. If the methylated version of the construct cannot enter the nucleus, then we should see a strong preference for the unmethylated version attached to a construct that has certainly entered the nucleus. This is not what we see, however: the methylated version of the construct is present on 1242 out of the 2677 reads which equates to 46%, in line with the proportion of methylated construct in the injected DNA (44.6%) (See Figure 4D). We therefore conclude that 6mA marks do indeed prevent excision of an IES in the developing macronucleus, at high efficiency.

**Figure 4.**
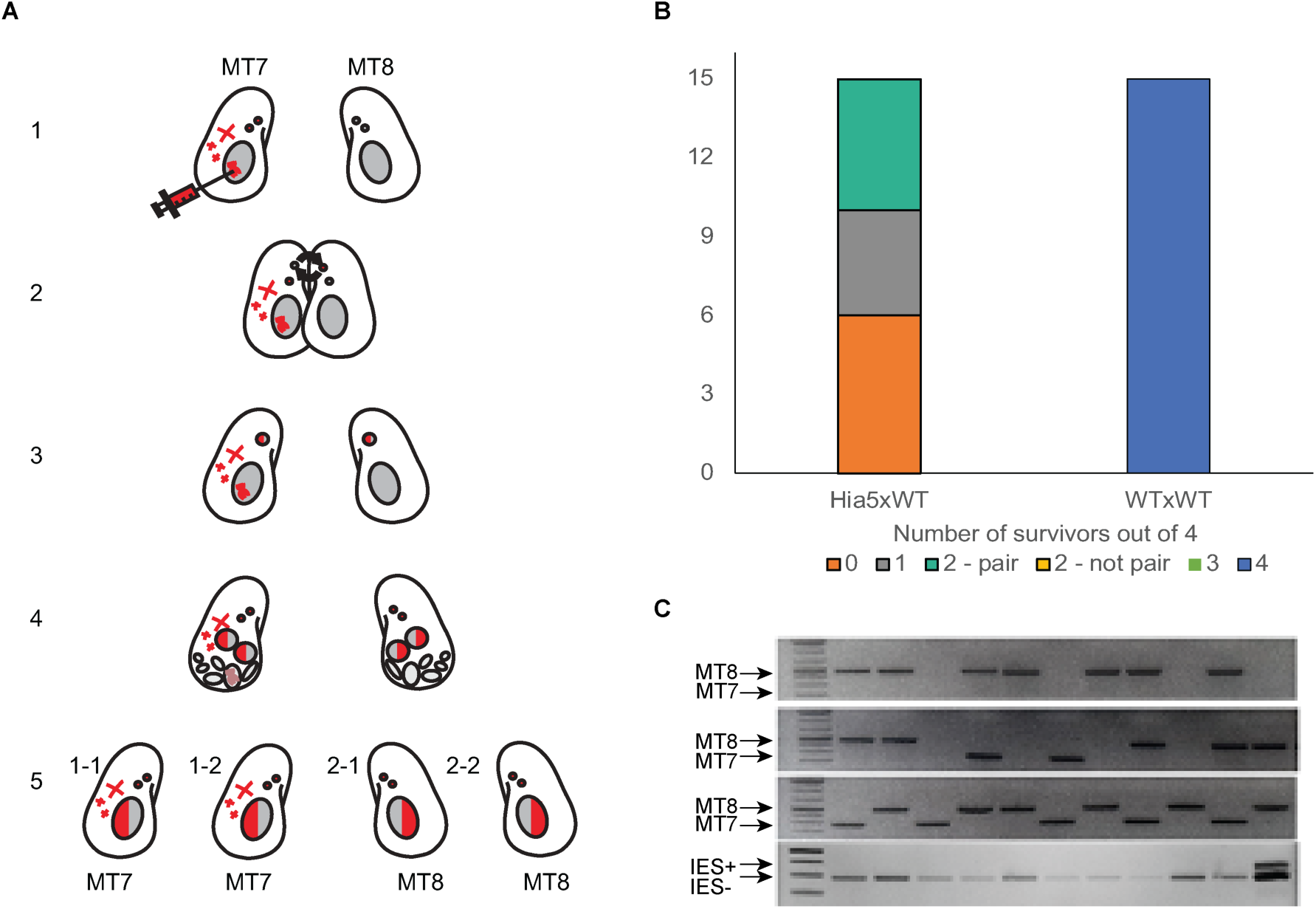
Conjugation experiment to assess whether the ectopic methylation phenotype seen in Hia5-expressing cells is transmitted through the MIC, as would be expected for a transgenerational epigenetic mark. (A) Schematic depicting the process of conjugation and the transmission of injected DNA, Hia5-dependent modifications and possible cytoplasmic or RNA modifications. 1) cells of mating type 7 (left) are injected into the MACs, and Hia5-CenH3 is transported to the MICs where it methylates DNA. Methylated MICs are red. Injected DNA in the MAC is represented by a red double helix, and possible cytoplasmic and/or RNA effects are represented by red crosses in the cytoplasm. Wild-type mating type 8 cells are depicted on the right, with grey DNA in MICs and MACs. 2) Cells conjugate and exchange one meiotic MIC, so both contain one modified and one unmodified MIC. 3) and 4) Haploid meiotic pronuclei fuse and then divide mitotically to produce two new MICs and two new developing MACs. The maternal MAC fragments. New MICs and new MACs now both carry Hia5-deposited methylation, shown in red. 5) Four karyonides from the conjugation are produced, each containing Hia5-deposited marks in both MIC and MAC. If cells 2-1 and 2-2 (mating type 8) have the same phenotype as cells 1-1 and 1-2 (mating type 7) then the phenotype was inherited from the germline MIC. (B) Survival test following Hia5 expression and conjugation, or autogamy. “Pair” means two cells survived originating from the same parent, “not pair” is two cells from opposite parent cells. (C) Mating type PCR or IES retention PCR of daughter cells after conjugation.

### 6mA is a transgenerational mark carried from germline to soma in Paramecium

While it is clear that 6mA blocks excision when placed on an IES, we still needed to ascertain that the mark is carried trans-generationally, from germline to daughter soma: from MIC to MAC. The Hia5 injection experiments demonstrate lethality and IES retention in progeny of cells where a generalised 6mA methyltransferase is targeted to MICs. However, despite being only visible in the MICs, we could not be certain that the Hia5 was not present in low quantities in the MAC and/or the cytoplasm, potentially methylating MAC DNA and/or RNA. The lethality and IES retention following Hia5 injection could be due to gene disruption or other indirect effects from Hia5 presence in the cells. To address this issue, we crossed Hia5-injected cells with WT cells of the opposite mating type. Paramecium has two mating types, termed 7 and 8, which will conjugate with each other under the correct conditions. We performed the Hia5 injections on mating type 7 cells, which we then conjugated with WT mating type 8 cells. The experiment is depicted in Figure 4A. As mentioned, following meiosis, two conjugating cells exchange a haploid meiotic pronucleus which then fuses with the pronucleus from the conjugating cell. The nuclei travel from their maternal cell to its conjugating partner via a cytoplasmic bridge. No MAC fragments pass across the bridge and negligible quantities of cytoplasm are exchanged. Therefore, if any effect from Hia5 expression is transmitted to the daughter cells, it must come from the meiotic MIC. We allowed cells to complete conjugation and then picked the daughter cells into separate wells to assess their viability, growth, and possible IES retention. Each exconjugant contains two developing MACs which each go to one daughter cell after the exconjugant undergoes karyonidal division. Therefore the offspring from one conjugating pair consists of four cells, two from each parental cell (see Figure 4A). If the offspring that arose from the WT cell display the same phenotype as those from the Hia5-expressing cell, then we can conclude that the phenotype was transmitted through the meiotic pronucleus. However, if the phenotype is not transmitted via the MIC but comes from indirect effects on e.g. MAC or cytoplasm arising from Hia5 presence in the maternal cell, then we would expect offspring deriving from the WT cell lineage not to have any abnormal phenotype. Results are shown in Figure 4B. 15 sets of cells were picked and scored for viability after four days. In 10 out of 15 pairs, at least one karyonid stemming from the WT conjugation partner died. This demonstrates that the lethality phenotype is transmitted via conjugation, and must therefore be carried in the MIC. We believe that the milder phenotype in the conjugated cells compared to autogamy is simply due to a dilution in the amount of aberrant methylation present, since a methylated Hia5 pronucleus fuses with a WT pronucleus with normal methylation patterns. Since *Paramecium* mating types cannot be distinguished visually, we performed PCR to assess to which mating type the surviving cells belong. Since we performed the Hia5 injections on mating type 7 cells, this would tell us whether surviving cells were preferentially the daughters of WT cells (mating type 8). As shown in Figure 4C, in the matings where two cells survived, all of the survivors were daughters of the WT mating type 8 partner. However, in the matings where three cells died, half the surviving single cells were daughters of the Hia5-expressing, mating type 7 cell. This discrepancy could be because the Hia5-expressing cells cannot perform conjugation as efficiently as WT cells, and in some cases conjugation is not completed. The result of a failure to complete nuclear exchange would be survival of the WT partner and its descendants and death of the Hia5-expressing partner, which is what we see in five cases. Nonetheless, in the majority of the cells tested, the lethality phenotype was transmitted through the MIC, demonstrating that the Hia5-mediated aberrant methylation in the MIC caused lethality in the next generation.

To assess IES retention in the progeny of conjugations between Hia5-expressing cells and WT cells, we performed IES retention PCR as before. We did not detect any IES+ bands in any of the cells, including the daughter cells of the Hia5-expressing cells themselves (Figure 4C). This could be explained by the Hia5-effect being weakened by dilution, as described above. The IESs may be retained at levels undetectable by PCR.

## DISCUSSION

We have demonstrated that 6mA is an epigenetic mark in Paramecium that is preferentially present on MDS, and which prevents IES excision on sequences where it is located. It is present in vegetative germline DNA, and remains there during sexual reproduction and development as IESs are excised and the genome is amplified. We believe that 6mA constitutes a transgenerational epigenetic mark in Paramecium, although mechanistic questions pertaining to its deposition, maintenance and how it exerts its protective effect still remain.

We do not know which DNA methyltransferase is responsible for depositing and/or maintaining 6mA marks on the Paramecium MIC. There are at least 14 Paramecium genes with putative 6mA methyltransferase functions^53^. During vegetative growth, Paramecium MICs divide mitotically with each cell division. To maintain 6mA marks, we expect that methylation marks are transferred to nascent DNA strands during or after replication, using the parental DNA methylation patterns as a template, similarly to Dnmt1 for cytosine methylation^54^. After meiosis, nuclear exchange and fusion, and the start of macronuclear development, 6mA is used as a protective mark to ensure that MDSs are not aberrantly excised, and to help the cellular excision machinery to distinguish MDS from IES. During this process, the genome is amplified up from 2n in the MIC to ∼800n in the MAC, while IES excision occurs concomitantly. Therefore, 6mA marks must be faithfully transferred to the amplifying MAC chromosomes if they are to exert their protective effect. We therefore anticipate that the expression profile of the methyltransferase in question will be low but consistent during vegetative growth, and should increase during the second half of development when the genome is being amplified. Further work will be required to determine which methyltransferase(s) are responsible for establishing and maintaining MIC 6mA marks.

According to our analyses, 6mA is present on approximately 0.6% of VATB tetranucleotides in MDSs. While a significant DNA modification, this is still overall a low level if we believe 6mA to be an important mark distinguishing MDS from IES. After all, in a genome consisting of 78% AT, only around 5% of tetranucleotides can be expected to be VATB. If only 2% of these are methylated with 6mA, 6mA marks will be present only in 0.1% of tetranucleotides. As described in the introduction, Paramecium IESs are ubiquitous in the genome and almost all of them exist within protein-coding genes, making their precise excision paramount for the survival of the cell. An epigenetic mark present at such low levels will not be sufficient to distinguish IES from MDS with any precision. How then can 6mA exert a meaningful effect during genome rearrangement? One possible explanation is that 6mA is deposited on particular sequences that otherwise could be “hot spots” for aberrant excision. It is known that several different factors are involved in helping the excision machinery to identify IESs, these include inverted repeats, chromatin marks and particular end sequences. It is also known that certain lengths of IES are strongly preferred over others; these follow a sinusoidal distribution with a period of 10.5 bp^25^. It is therefore feasible that certain loci within the genome are more prone to being incorrectly excised, and that 6mA modifications have evolved to help ensure that this does not occur.

Another possibility is that the protective effect that 6mA marks exert may be far-reaching, and a 6mA modification signals that all surrounding DNA within a certain area must be retained, or all DNA up until another, opposing signal (such as an IES boundary) is encountered. In this scenario, the’off’ signal limiting the reach of the 6mA protection would be as important as the 6mA mark itself, implying that 6mA protection evolved as a part of a belt-and-braces system to ensure correct genome rearrangement.

Further work will be required to establish the mechanism of 6mA genome protection.

One interesting observation made during this study is that Paramecium 6mA is preferentially located on VATB tetranucleotides. This preference is also observed in Oxytricha and Chlamydomonas, two evolutionarily distant protozoans, but not in Tetrahymena, a more closely related ciliate to Paramecium. This raises the question of whether the VATB preference predates the divergence of algae and ciliates and was lost in the Tetrahymena lineage, or whether it is an example of convergent evolution, and for some reason VATB has an advantage as a location for 6mA modification. Identification of the methyltransferases in question may help to shed light on this question. As mentioned, avoidance of TATA in Paramecium makes sense if IES end sequences are actively avoided as targets for 6mA, which is consistent with a protective role in genome rearrangement. This preference may not exist in Tetrahymena, where IES ends have a TTAA consensus, and where it is not known if 6mA has a role in genome rearrangement. The protective role of 6mA in Paramecium may have evolved recently; Paramecium has undergone two whole-genome duplications since its divergence from Tetrahymena^55^.

Whole-genome duplications give the genome a degree of plasticity in allowing the evolution of new functions for genes.

## STAR★METHODS

### KEY RESOURCES TABLE

| REAGENT OR RESOURCE | SOURCE | IDENTIFIER |
| --- | --- | --- |
| Bacterial and virus strains |  |  |
| Endura competent *E. Coli* cells | Lucigen | Cat #: 60242-0 |
| Feeding bacteria HT115 *E. Coli* cells | Gift from Eric Meyer (ENS, Paris) | HT115 |
| *Klebsiella pneumoniae* non-virulent strain, food source for *Paramecium* | Gift from Eric Meyer (ENS, Paris) | N/A |
| Chemicals, peptides, and recombinant proteins |  |  |
| Wheat Grass Powder | Pines International, Lawrence, KS | N/A |
| B-sitosterol | Calbiochem, Millipore | Cat #: 567152; CAS: 83-46-5 |
| Acetonitrile | Millipore | Cat #: 75-05-8 |
| Formic Acid | Millipore | Cat #: 64-18-6 |
| Benzonase | Millipore | Cat #: 70664 |
| Phosphodiesterase I | Millipore | Cat #: P3243 |
| Alkaline Phosphatase | Millipore | Cat #: 524572-5KU |
| Proteinase K | Promega | Cat #: MC5005 |
| T4 DNA Ligase | New England Biolabs | Cat #: M0202 |
| Tri Reagent | Sigma-Aldrich | Cat #: T9424 |
| Critical commercial assays |  |  |
| Q5 Site-Directed Mutagenesis Kit | NEB | Cat #: E0554S |
| GoTaq G2 DNA polymerase | Promega | Cat #: M7848 |
| Phusion High-Fidelity DNA Polymerase | Thermo Scientific | Cat #: F-530L |
| QIAGEN Plasmid Midi Kit | QIAGEN | Cat #: 12143 |
| pGem-T Easy Vector System | Promega | Cat #: A3600 |
| Wizard® SV Gel and PCR Clean-Up System | Promega | Cat #: A9281 |
| Deposited data |  |  |
| Mass spectrometry | This paper |  |
| *Paramecium tetraurelia* SMRT-seq | This paper | NCBI: PRJNA1373137 |
| *Paramecium tetraurelia*, all Illumina data (gDNA-seq, Small RNA-seq, Amplicon-seq) | This paper | NCBI: PRJNA1373137 |
| Experimental models: Organisms/strains |  |  |
| *Paramecium tetraurelia*, strain 51 | Gift from Eric Meyer (ENS, Paris) | N/A |
| Oligonucleotides |  |  |
| See Table S1 for complete list of oligonucleotides | This paper | N/A |
| Recombinant DNA |  |  |
| CenH3-GFP-Hia5 | This paper | N/A |
| CenH3-Hia5 | This paper | N/A |
| CenH3-Hia5-D194A | This paper | N/A |
| Dcl5 silencing construct | Sandoval et al., 2014 | N/A |
| PGM silencing construct | Baudry et al., 2009 | N/A |
| Software and algorithms |  |  |
| Geneious 2025 | http://www.geneious.com/ | N/A |
| WebLogo 3 | WebLogo, Crooks et al., 2004 | http://weblogo.threeplusone.com/ |
| Other |  |  |
| Amicon Ultra-2 Centrifugation Filter Unit | Millipore | Cat #: UFC200324 |
| Ultrafree-MC Centrifugal Filter | Millipore | Cat #: UFC30GV25 |
| Amicon® Ultra Centrifugal Filter | Millipore | Cat #: UFC5030 |

### Contact for Reagent and Resource Sharing

Further information and requests for resources and reagents should be directed to and will be fulfilled by the Lead Contact, Mariusz Nowacki.

### EXPERIMENTAL MODEL AND STUDY PARTICIPANT DETAILS

All experiments were carried out using *Paramecium tetraurelia* strain 51 (mating type 7 unless otherwise specified), cultured at 27 °C in wheat grass powder (WGP; Pines International, Lawrence, KS) medium bacterized with *Klebsiella pneumoniae* and supplemented with 0.8 mg/mL β-sitosterol. Autogamy was induced by starvation.

## METHOD DETAILS

Mass spectrometry analysis of DNA modifications Macronuclear DNA (see Macronuclear DNA extraction and Sequencing, below) was purified from PGM-KD and empty vector silencing cultures at 2 days post autogamy. Up to 10 µg of sample DNA was hydrolyzed to nucleosides in the following enzymatic digestion process: The samples were added to a mixture containing 0.2 u Benzonase (MilliporeSigma, 70664), 0.5 mU of Phosphodiesterase I from *Crotalus adamanteus* venom (MilliporeSigma, P3243) and 0.5 U of Alkaline Phosphatase from calf intestine (MilliporeSigma, 524572-5KU) for 2 h at 37°C in a final volume of 100 µl of digestion buffer (5 mM TRIS pH 8.0, 20 mM NaCl and 1 mM MgCl_2_). As an internal loading control, an equal volume of ^13^C, ^15^N-labelled uridine (MilliporeSigma, 645672) in 0.2% formic acid was added to the samples. These were then filtered through 30 kDa Molecular Weight Cut-Off filters (Amicon® Ultra Centrifugal Filter, 30 kDa MWCO, MilliporeSigma, UFC5030) to remove the proteins. 2 μL of each sample was injected in triplicate for LC-MS/MS analysis.

### DNA Nucleoside Mass Spectrometry

#### Nucleoside Standards and Chemical Products

Nucleoside standards were purchased from MillioporeSigma and Biosynth (formerly Carbosynth, Compton, United Kingdom). All chemicals purchased were grade LC-MS: Acetonitrile (1.00029, CAS: 75-05-8), water (1.15333, CAS: 7732-18-5), and formic acid (5.33002, CAS: 64-18-6) were purchased from MilliporeSigma.

#### UPLC-MS Method

Samples were resolved over 15 minutes through an Acquity 100 × 2.1 mm C-18 HSS T3 column using a Thermo Scientific U3000 UPLC system on a gradient of 2–98% (0.1% formic acid/acetonitrile) and analysed on a QExactive-HF Orbitrap High Resolution Mass Spectrometer (Thermo Fisher Scientific, IQLAAEGAAPFALGMBFZ) in positive full-scan mode. Initial conditions were regenerated by rinsing with 100% solvent A (0.1% formic acid, 2% Acetonitrile in water) for an additional 2 min. The flow rate was 0.2 mL/min, and the column temperature was 25°C.

The results were deconvoluted using the accompanying Xcalibur Software (Thermo Fisher Scientific). Nucleosides were assigned based on retention time and exact mass, and quantified relative to calibration curves generated from pure standards. Modified nucleosides were expressed as a percentage of their precursor nucleoside (ie N6mdA as a percentage of dA).

| Nucleoside | Molecular Formula (M) | Accurate [M+H]<sup>+</sup> Mass (Da) | Retention Time (min) |
| --- | --- | --- | --- |
| deoxycytidine | C9H13N3O4 | 228.0984 | 1.96 |
| deoxyadenosine | C10H13N5O3 | 252.1097 | 4.8 |
| deoxyguanosine | C10H13N5O4 | 268.1046 | 6.11 |
| thymidine | C10H14N2O5 | 243.0981 | 7.5 |
| deoxy5methylcytidine (5-mdC) | C10H15N3O4 | 242.1141 | 3.42 |
| deoxy5hydroxymethylcytidine (5-hmdC) | C10H15N3O5 | 258.109 | 1.88 |
| deoxy5formylcytidine (5-fdC) | C10H13N3O5 | 256.0933 | 6.09 |
| N6methyldeoxyadenosine (N6-mdA) | C11H15N5O3 | 266.1253 | 8.04 |
| 1-methyldeoxyguanosine (1-mdG) | C11H15N5O4 | 282.1202 | 8.38 |
| heavy_uridine (Internal Standard)* | (13C)9H12(15N)2O6 | 256.1198 | 3.02 |

#### Construction of CenH3-Hia5 fusions and microinjection

The coding sequence of the non-specific N⁶-adenine methyltransferase Hia5 (*Haemophilus influenzae*, GenBank: AAV37176.1) was codon-optimized for *Paramecium tetraurelia*, synthesised (GenScript), and fused to the C-terminus of the centromeric histone variant CenH3 (PTET.51.1.G4660001), along with a C-terminal GFP tag. Constructs were cloned between the endogenous 5′ and 3′ untranslated regions (UTRs) of the CenH3 gene into the pGEM-T vector (Promega). A non-GFP variant was generated by removing the GFP tag, and a catalytically inactive (D194A) mutant was produced via site-directed mutagenesis (Q5 Site-Directed Mutagenesis Kit, New England Biolabs). Linearized plasmids (5-10 µg) were microinjected into the somatic macronucleus of vegetative cells using a FemtoJet microinjector (Eppendorf) as described^56^.Successful transformants were identified by GFP fluorescence or PCR genotyping.

#### GFP localization and imaging

Following microinjection, cells were collected, washed twice in 10 mM Tris-HCl (pH 7.4), fixed in 70% ethanol, and stored at 4 °C. For imaging, cells were washed twice with 1× PBS (pH 7.4) and stained with DAPI. Imaging was performed using a phase-contrast inverted microscope (Axiovert A1, Zeiss), and images were processed with ZEN 2 software (Zeiss).

#### Macronuclear DNA extraction, sequencing and IES retention PCR

High-quality macronuclear DNA was isolated using a modified sucrose gradient ultracentrifugation protocol^5^. Briefly, Paramecium cells were lysed in a non-ionic detergent buffer and mechanically homogenized on ice to release macronuclei, which were purified via sucrose cushion ultracentrifugation (35,000 rpm, 1 h, 4°C; SW40Ti rotor). Purified macronuclei were lysed in SDS/proteinase K buffer and incubated overnight at 55°C. DNA was extracted using phenol:chloroform:isoamyl alcohol, followed by dialysis against Tris-EDTA buffer. The recovered DNA was concentrated using centrifugal filter units (30 kDa MWCO; Amicon Ultra-2, Millipore), quantified by spectrophotometry, and quality-checked by agarose gel electrophoresis.

Sequencing libraries were made using TruSeq Nano DNA High Throughput Library Prep Kit (96 samples, illumina, 20015965) in combination with IDT for Illumina-TruSeq DNA UD Indexes v2 (96 Indexes, 96 Samples, illumina, 20040870) according to the kit reference guide (illumina, Part # 15041110 Rev. D June 2015). Pooled DNA libraries were sequenced paired-end on a shared NextSeq1000 P2 Reagent Kit (300 cycles; illumina, 20100985) on an illumina NextSeq1000. The quality of the sequencing run was assessed using illumina Sequencing Analysis Viewer (illumina version 2.4.7) and all base call files were demultiplexed and converted into FASTQ files using illumina bcl2fastq conversion software v2.20. All steps from DNA QC to sequencing data generation and data utility were performed at the Next Generation Sequencing Platform, University of Bern, Switzerland. IES retention scores were calculated with the MIRET component of ParTIES^57^. IES retention PCR was performed on a genomic region flanking a representative internal eliminated sequence (IES), using primers listed in Table S1.

#### Survival test

Autogamy was induced by starvation. For the survival assay, 30 post-autogamous cells were individually transferred to three-well glass slides containing medium inoculated with avirulent *Klebsiella pneumoniae*. Cells were monitored over three days (corresponding to approximately 12 cell divisions) and categorized into three phenotypic groups.

#### Sequencing of Small RNA

Small RNA sequencing was performed by Fasteris SA (Geneva, Switzerland). Approximately 600,000 Paramecium cells were harvested, washed twice in 10 mM Tris (pH 7.4) and cell pellets were snap-frozen in liquid nitrogen. Total RNA was extracted using TRIzol reagent (Sigma-Aldrich) according to the manufacturer’s protocol. Small RNA species were enriched by polyacrylamide gel electrophoresis, and RNA fragments of 17–35 nt were excised and recovered. Libraries were prepared from 3 µg total RNA using the Illumina TruSeq Small RNA kit (adapter ligation, RT, and indexed PCR), purified using CRL/HRL reagents, and quality-controlled on an Agilent DNA 1000 system. Sequencing was performed on an Illumina NextSeq instrument using single-end 1 × 75 bp high-output mode. Adapter trimming and UMI extraction were performed using UMI-tools v0.2.3. The following alignments were performed by Hisat2^58^ against (the genome of *Klebsiella*, L4440 plasmid, *P. tetraurelia* rRNA, mitochondrial genome, xxx, MAC, MAC+IES, MIC), respectively.

#### Silencing Specific Genes by dsRNA Feeding

PGM and DCL5 were silenced by feeding *Paramecium* cells with dsRNA-producing *E. coli*, following a modified protocol as previously described^56^. Bacterial cultures containing the RNAi constructs were grown overnight in LB with ampicillin (0.1 mg/mL) and tetracycline (0.0125 mg/mL), diluted 1:100 in 1× WGP with ampicillin, and incubated for 7–8 h at 37 °C. After a 1:5 dilution in fresh WGP, cultures were induced with 0.4 mM IPTG overnight. The culture was cooled to 27 °C, supplemented with β-sitosterol (0.8 mg/L), and used to feed Paramecium at a density of 200 cells/ml. Autogamy was induced by starvation. Efficient silencing of PGM and Dcl5 was confirmed by PCR analysis of retained IESs, whose excision is specifically dependent on the respective gene.

#### Synthetic MDS-IES-MDS constructs and injection during autogamy

A 30-bp IES (IESPGM.PTET51.1.160.128593 in Paramecium Database) along with 10bp upstream and downstream flanking sequence was synthesised (Twist Bioscience) with two 30bp of flanking MAC sequence from other genomic regions on each side. Four ApT sites within the IES were chemically modified to contain N⁶-methyladenine on both strands (custom synthesis, Genscript). A 3-bp barcode distinguished methylated and unmethylated versions (CCC and ATG respectively). Equal amounts were mixed and injected into the cytoplasm of Dcl5-silenced cells at mid-late autogamy stage (15-20h after starvation) as described^56^.

#### DNA extraction and PCR amplification for Synthetic MDS-IES-MDS constructs

Cells injected with synthetic MDS–IES–MDS constructs were allowed to complete development and grown for ∼10 vegetative divisions before genomic DNA extraction. Approximately 50,000 cells were harvested and DNA was isolated using the GeneElute Mammalian Genomic DNA Miniprep Kit (Sigma-Aldrich) according to the manufacturer’s protocol. A flanking primer pair spanning the synthetic construct (Table S1) was used for PCR amplification using Phusion High-Fidelity DNA Polymerase (Thermo Scientific, F-530L).

#### Amplicon sequencing and data analysis

The quantity, purity, and integrity of the PCR products were assessed using a Qubit 4.0 fluorometer (dsDNA HS/BR Assay Kit; Thermo Fisher Scientific), a DeNovix DS-11 FX spectrophotometer, and the Agilent FEMTO Pulse System (Genomic DNA 165 kb Kit), respectively. Sequencing libraries were generated using the NEBNext Ultra II FS DNA Library Prep Kit for Illumina (NEB, E7805L) with Unique Dual Index UMI Adaptors (NEB, E7395L), following the manufacturer’s instructions (Version 4.0_7/23), using 50 ng input DNA and 5 PCR indexing cycles. Library quality was evaluated with a Qubit dsDNA HS Assay (Thermo Fisher Scientific) and an Agilent Fragment Analyzer (HS NGS Fragment Kit). Pooled libraries were sequenced in paired-end mode (151:20:8:151) using NextSeq1000 P2 Reagent Kit (300 cycles) on a NextSeq1000 instrument and using MiSeq i100 Series 5M Reagent Kit (300 cycles) on a MiSeq i100 Plus instrument (Illumina). All DNA QC and sequencing were performed at the Next Generation Sequencing Platform, University of Bern, Switzerland. The data was analysed by Geneious Prime (2025).

#### Single-molecule real-time (SMRT) sequencing and 6mA detection

High-molecular-weight genomic DNA was extracted following the protocol optimized for long-read sequencing as previously described^59^. Briefly, cells were lysed in a preheated buffer containing SDS, EDTA, and polyvinylpyrrolidone, supplemented with RNase A. Protein and polysaccharide contaminants were precipitated using 5 M potassium acetate at 4°C. The lysate was clarified by centrifugation, and the supernatant was purified using magnetic carboxylate-modified beads (Serapure) with polyethylene glycol (PEG 8000)–based binding buffer. DNA-bound beads were washed twice with ethanol-based wash solution, air-dried, and eluted in EB buffer at 50°C. DNA purity and concentration were assessed via spectrophotometry and fluorometry, and integrity was verified by pulsed-field gel electrophoresis.

Library preparation and sequencing were performed at the Next Generation Sequencing (NGS) Platform, University of Bern. DNA extracted from vegetative cells was sequenced using the PacBio Sequel II system, and CCS reads (HiFi reads) were generated using CCS (v6.3.0). DNA from PGM-silenced and CenH3-Hia5 expressing cells was sequenced using the PacBio Revio system, with CCS reads generated using CCS (v8.2.0).

Alignments and kinetics feature calling were performed using SMRT Link (v25.2). Genome assemblies (MAC, MIC, and MAC+IES) were obtained from ParameciumDB (https://paramecium.i2bc.paris-saclay.fr). Modification thresholds were established based on the distribution of sequencing depth and inter-pulse duration ratios (IPDr). For vegetative data, bases with depth > 80x and log2(IPDr) > 1.5 were defined as modified. For the PGM-silenced and CenH3-Hia5 datasets, reads were classified using custom Perl scripts prior to modification identification: “IES++” (all IESs covered by the read were retained). Due to subsampling, the modification thresholds for these groups were adjusted to coverage > 20x and log2(IPDr) >1.

To identify MIC-derived reads in vegetative data, all reads were queried against a dataset of ∼45,000 IESs using BLAST^60^. A total of 4,014 reads showing > 98% identity to any IES were selected. Modified bases on these MIC reads were predicted using Fibertools^61^. Vegetative MIC reads was used for Sequence logos were generated using WebLogo3^62^. GC content used for the analysis was 27.74%, with the background composition set to 30% GC to reflect the genomic context of Paramecium.

#### Conjugation progress analysis and viability test

MT7 cells injected with the Hia5–CenH3 construct were screened by PCR to identify successfully transformed lines. Wild-type MT8 cells and Hia5-injected MT7 cells were cultured separately in 4 mL of 0.2× WGP medium bacterized with *Klebsiella pneumoniae* and incubated overnight at 27 °C. The following day, cells that had concentrated near the surface (∼1.5 mL) were collected from each culture and mixed to initiate conjugation as described^63^.

Individual mating pairs were isolated into fresh bacterized 0.2× WGP medium and monitored until pair separation. After separation, two ex-conjugant cells from each pair were transferred individually into three-well glass slides. Cells were cultured and monitored for three days, corresponding to approximately 12 vegetative divisions. Mating types of ex-conjugants were determined by PCR using established mating type-specific primers.

#### Data and Software Availability

All sequencing data have been deposited in NCBI under the BioProject accession number PRJNA1373137.

## QUANTIFICATION AND STATISTICAL ANALYSIS

All statistical analyses were performed in Microsoft Excel, as described in the respective figure legends.

## ACKNOWLEDGMENTS

We would like to thank Nasikhat Stahlberger for technical support.

## AUTHOR CONTRIBUTIONS

Conceptualization, S.A, and M.N.; methodology, X.L., S.A., A.H., L.L., and M.N.; Investigation, X.L., S.A., L.L., A.H., S.B., and M.N; writing—original draft, S.A.; writing—review & editing, X.L., C.E. and M.N.; funding acquisition, M.N,; resources, M.N. and C.K. supervision, S.A., and M.N:

## DECLARATION OF INTERESTS

No conflict of interests declared.

## DECLARATION OF GENERATIVE AI AND AI-ASSISTED TECHNOLOGIES

No AI-assisted tools used.

## SUPPLEMENTAL INFORMATION

Document S1. Figures S1-S2, Tables S1

**Figure S1.**
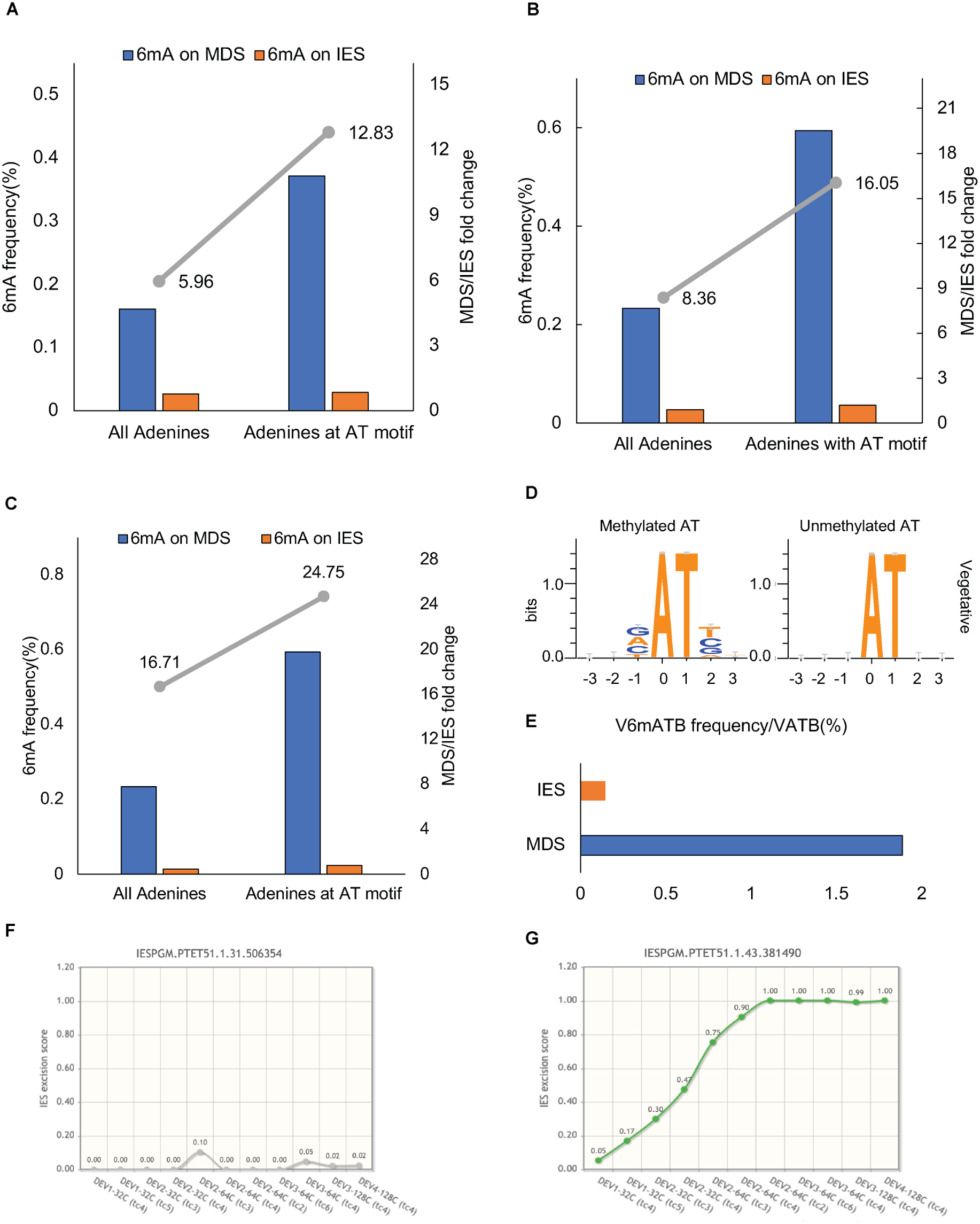
Further analysis of 6mA on MDS versus IES, including a wider dataset and removal of mis-annotated IESs. (A) 6mA levels on MDS versus IES, as in figure 1B after removal of mis-annotated IESs in the reference dataset. (B) vegetative MIC DNA reads were selected by including reads containing at least one IESs. 6mA frequencies are shown for all adenines, and for adenines within an AT motif. Fold change between MDS versus IES is indicated by the grey bar and the Y axis on the right. (C) 6mA levels on MDS versus IES, as in figure S1B after removal of mis-annotated IESs in the reference dataset. (E) 6mA frequency on MDS versus IES when only VATB motifs are taken into account from the PGM-KD DNA. (F) A representative mis-annotated IES showing excision score over development. The score is always at 0 and does not change, indicating that the IES is not excised during development. (G) A representative real IES for comparison, showing the excision score changing over time.

**Figure S2.**
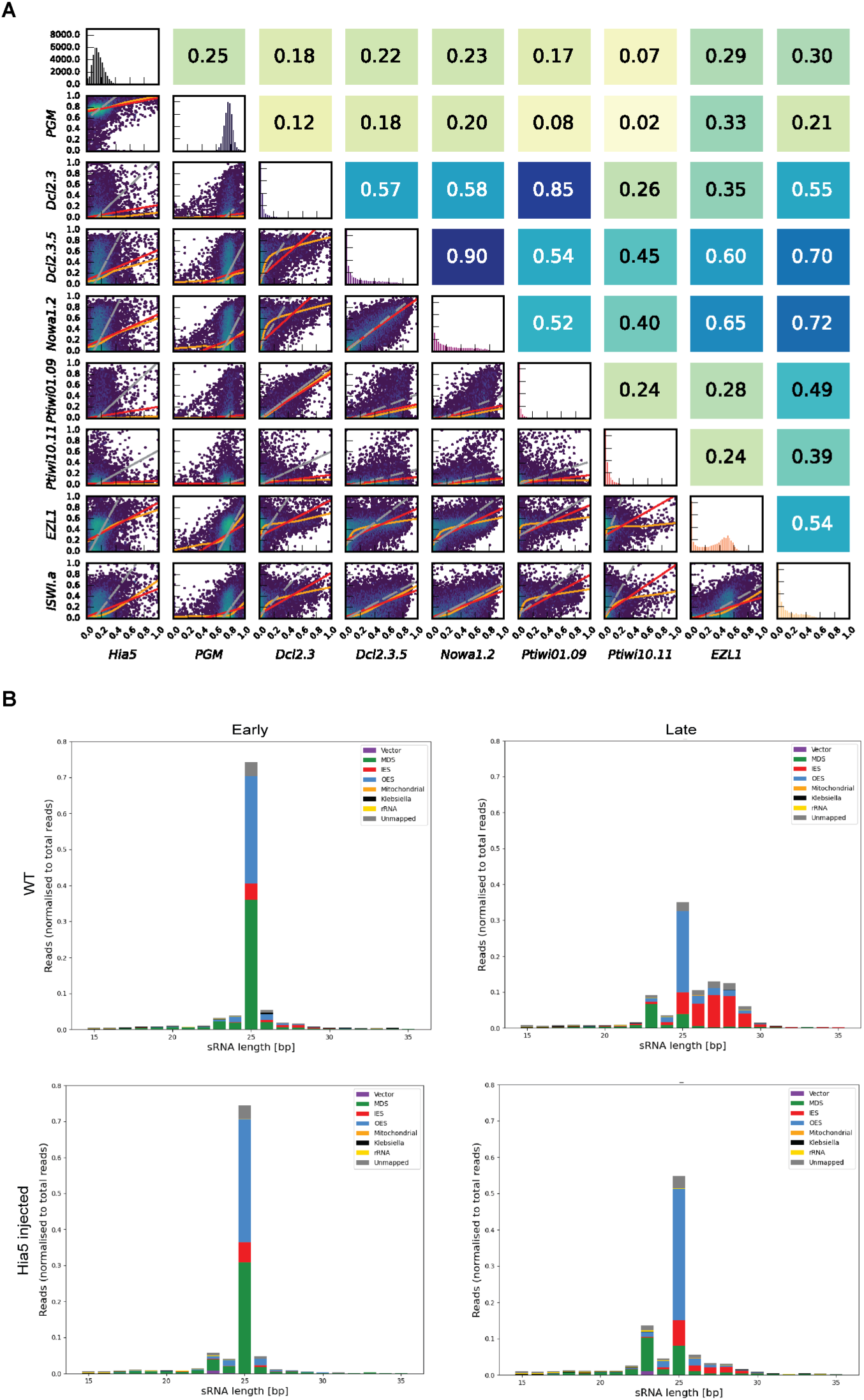
Assessment of whether Hia5 expression perturbs genome rearrangement through gene misregulation or small RNA pathways. (A) Correlation plots for IRS in Hia5-expression versus knockdowns of known genes: PiggyMac (PGM), Dcl2 and 3 (produce scanRNAs), Dcl2, 3 and 5 (all sRNAs in genome rearrangement), Nowa1 and 2 (RNA scanning), Ptiwi01 and 09 (Piwis that bind scnRNAs), Ptiwi10 and 11 (Piwis that bind iesRNAs), EZL1 (part of the PRC complex, methylates H3K9 and H3K27 during development). Correlation plots are generated as described^26^. Spearman’s rank correlation coefficient (rs) is given in the corresponding position diagonally opposite in the upper triangular matrix. Hia5 does not correlate with any of the analysed genes. (B) Small RNA distributions during development. The upper panels show wild-type autogamy with early versus late development. 25 nt scnRNAs are processed and the proportion of MAC-matching scnRNAs decreases over time. In later timepoints, iesRNAs appear which are produced from excised IESs, and therefore map to IESs. The lower panel shows sRNAs from Hia5-expressing cells. scnRNAs are processed normally, but iesRNAs are less abundant. Since IESs are retained in Hia5-expressing cells, there is less template for the production of iesRNAs and therefore their levels are lower. Otherwise, there is no defect in the sRNA pathway.

---

**Supplementary Table S1.**
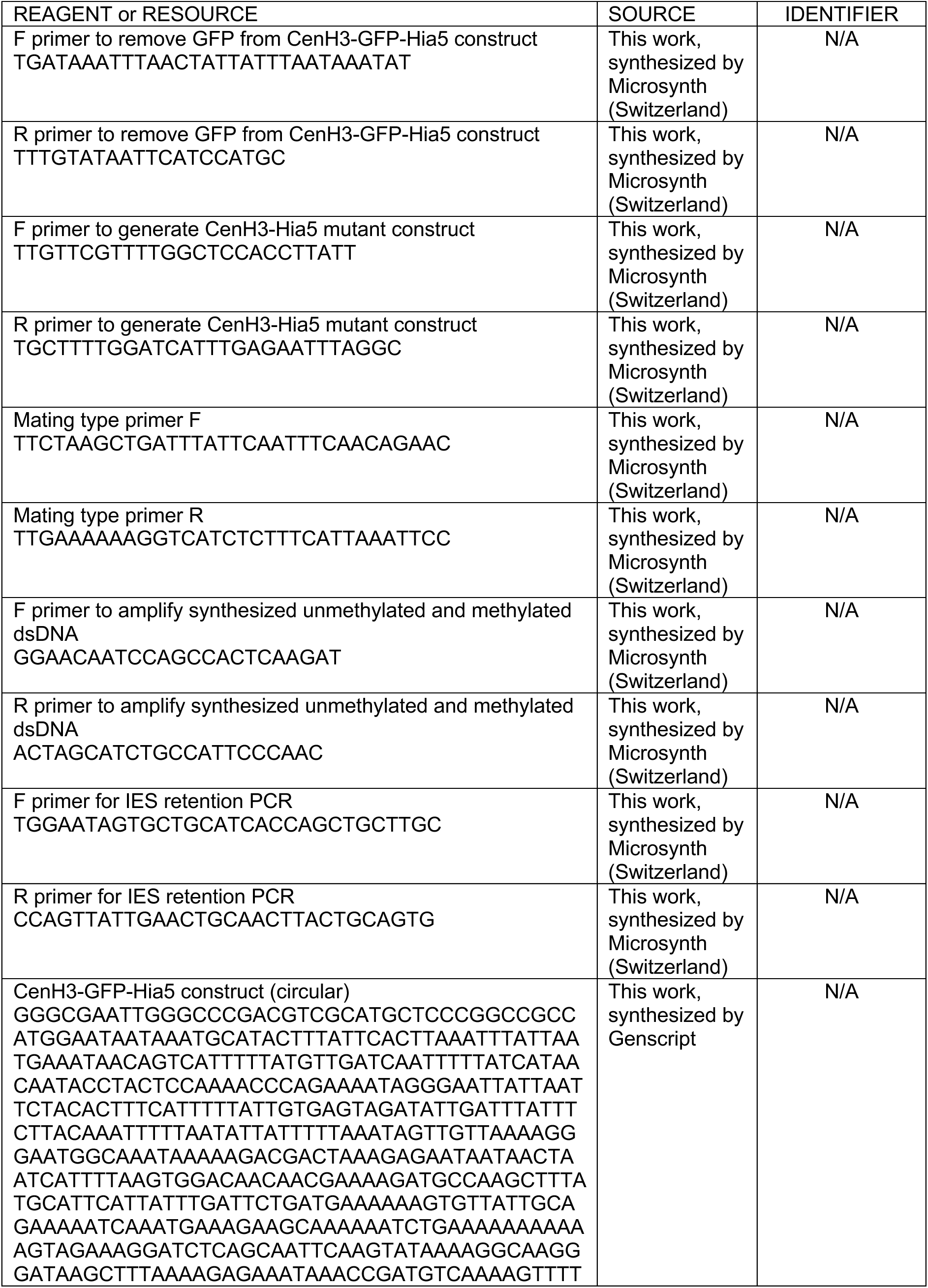
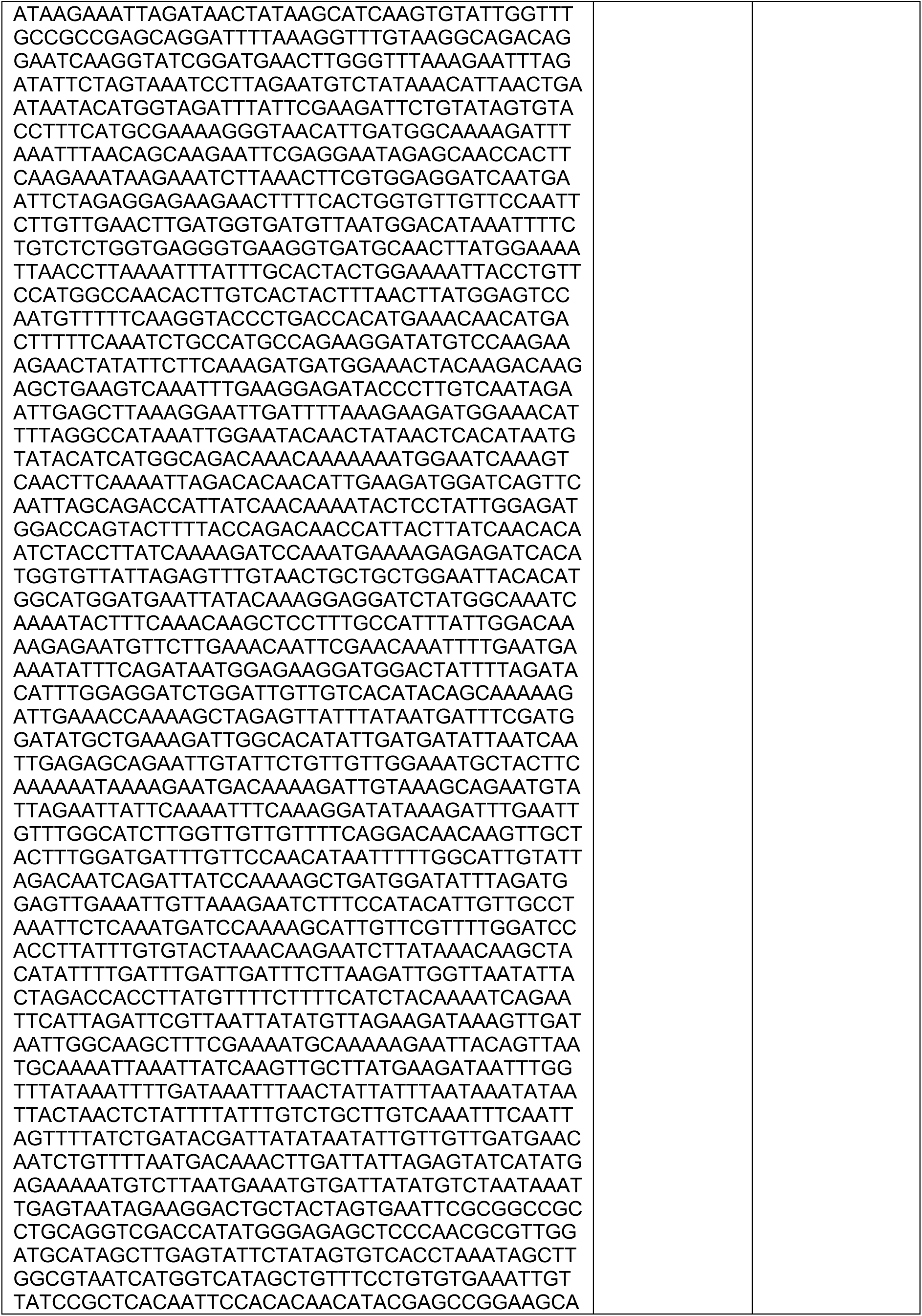
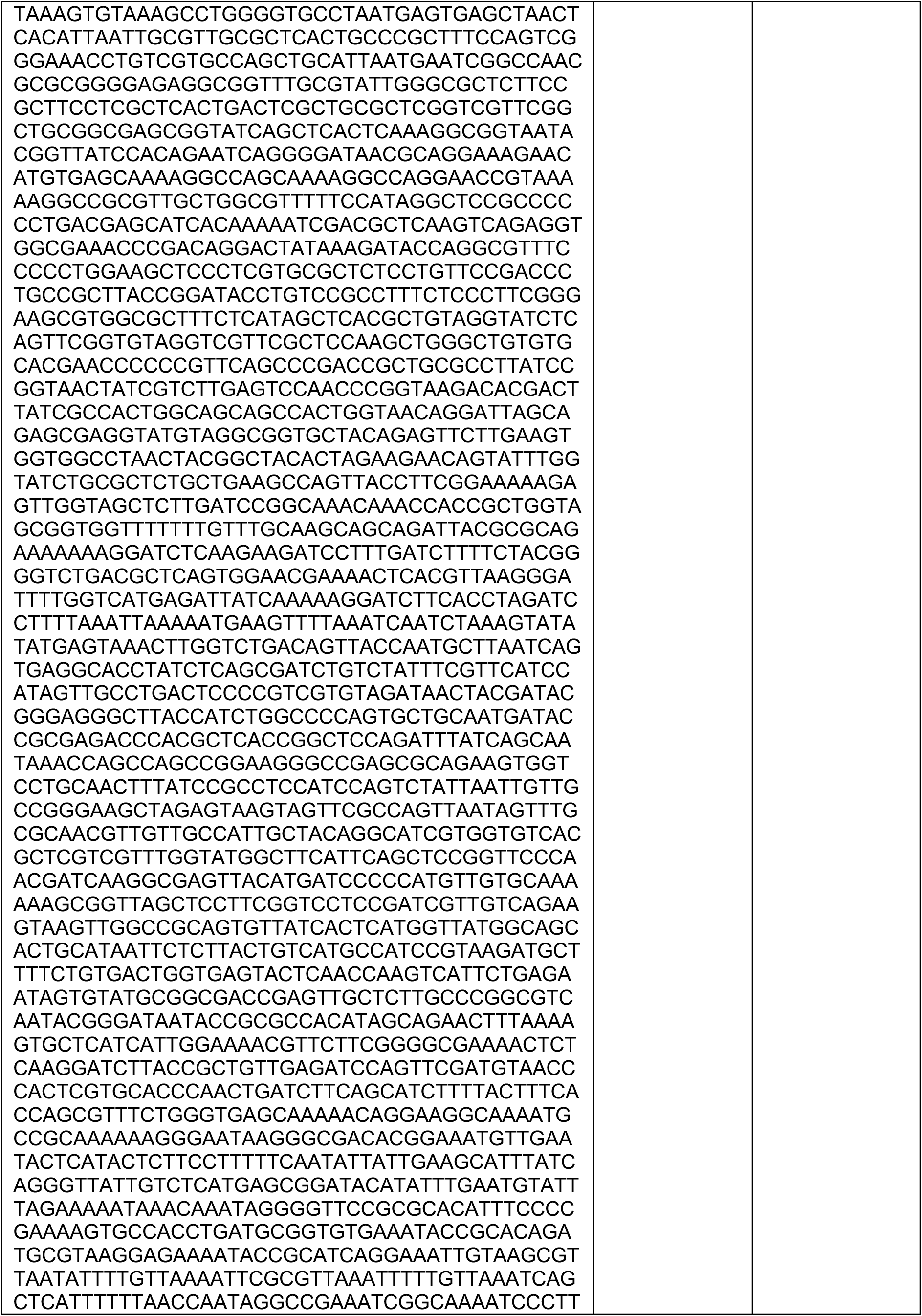
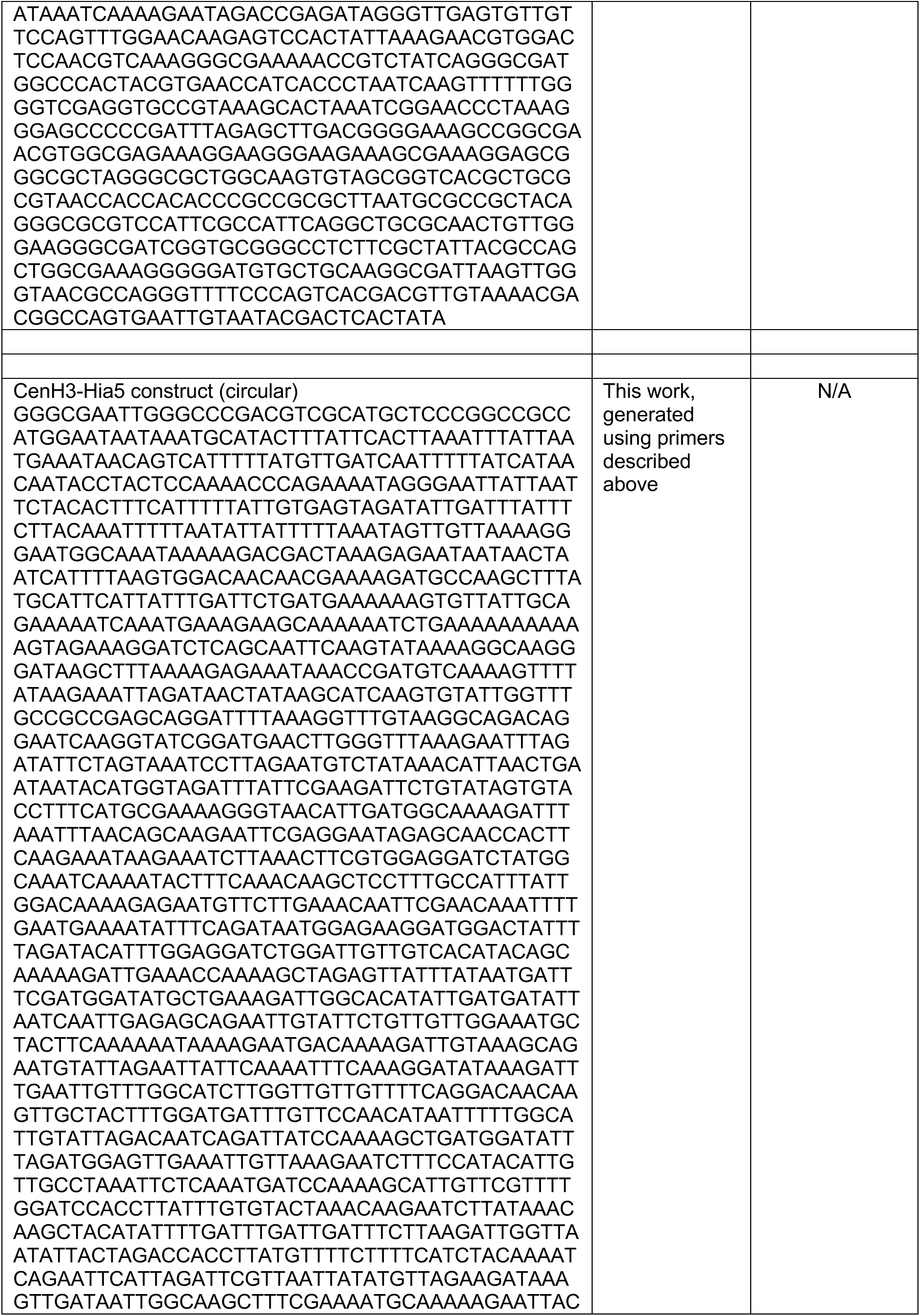
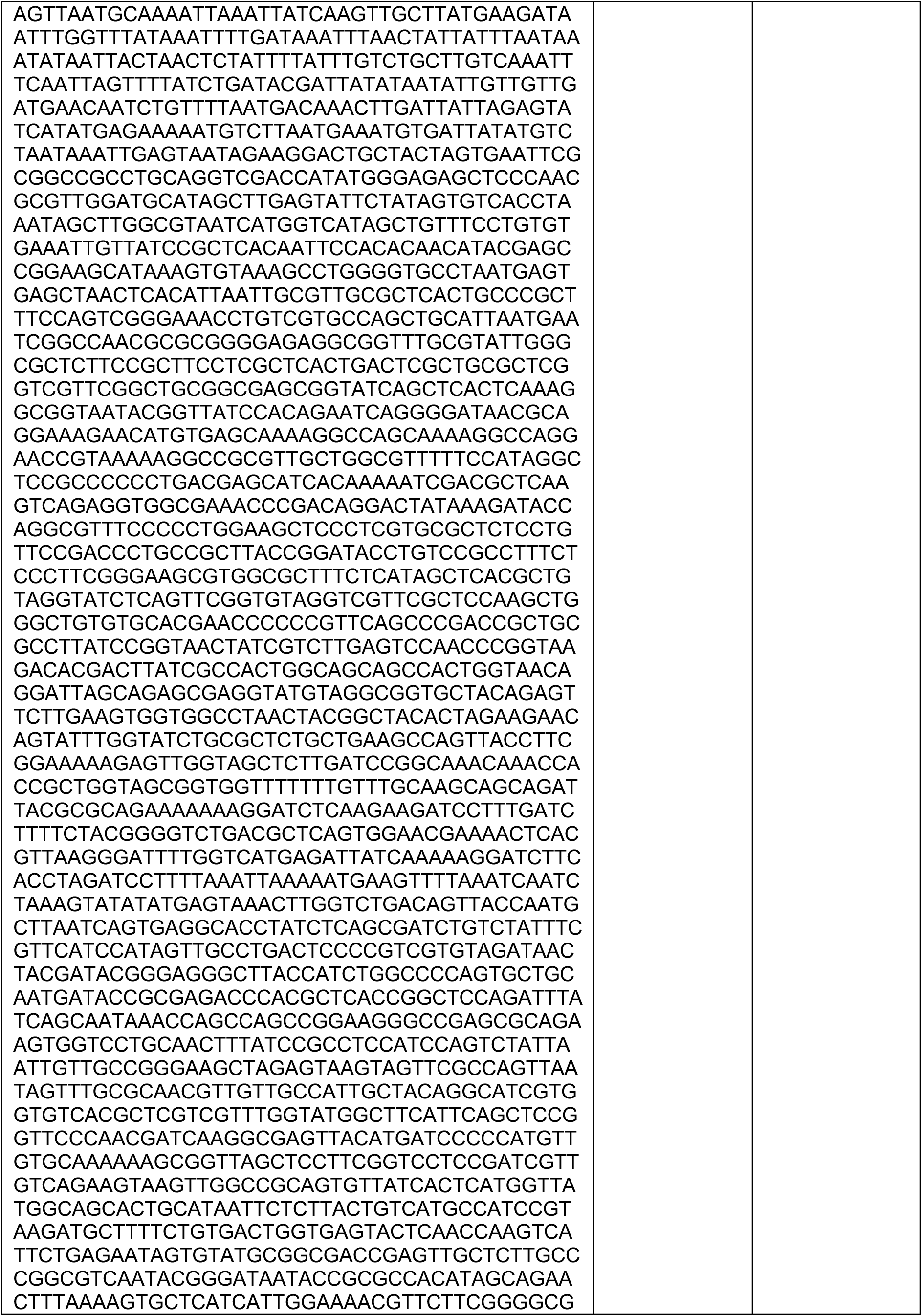
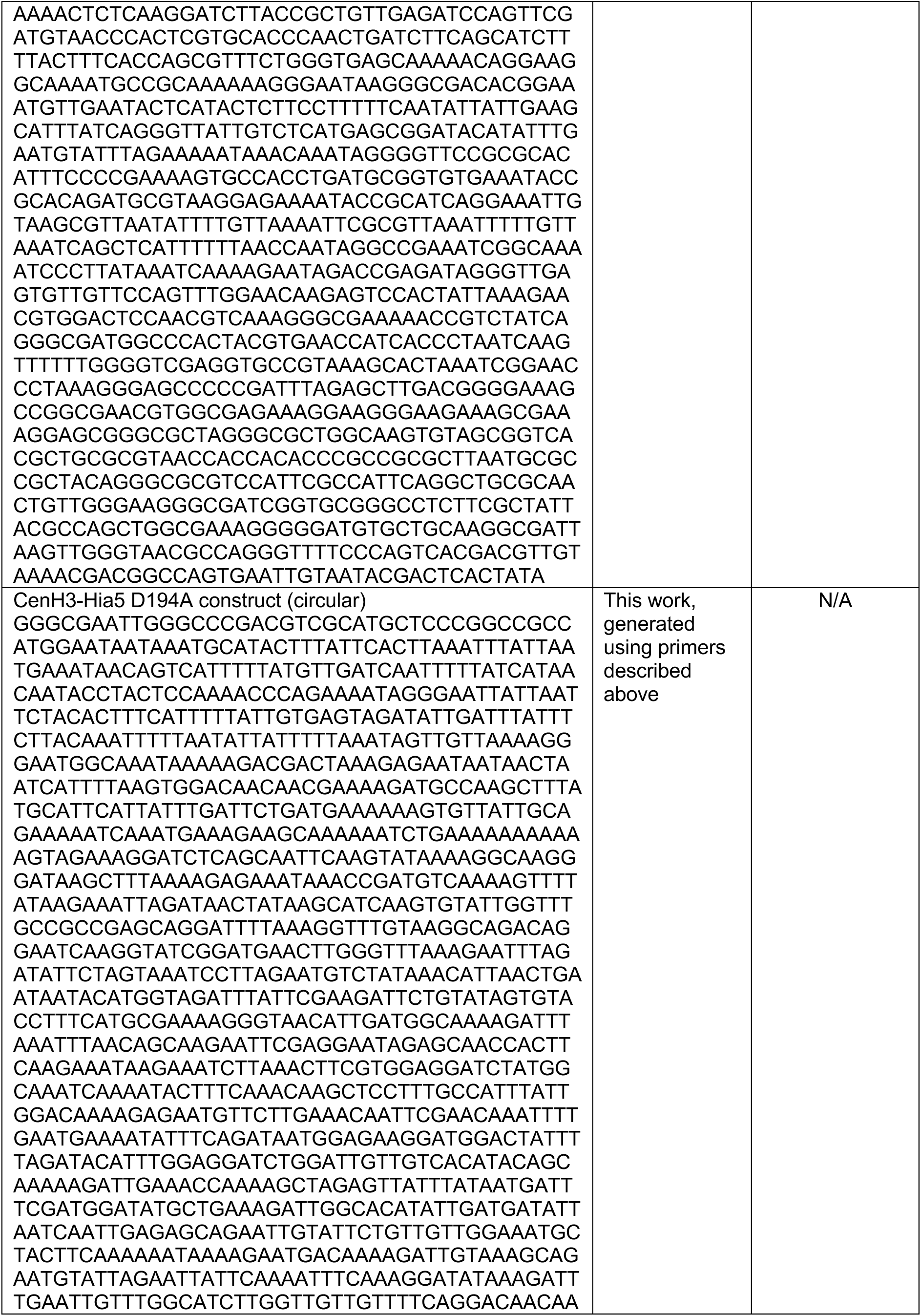
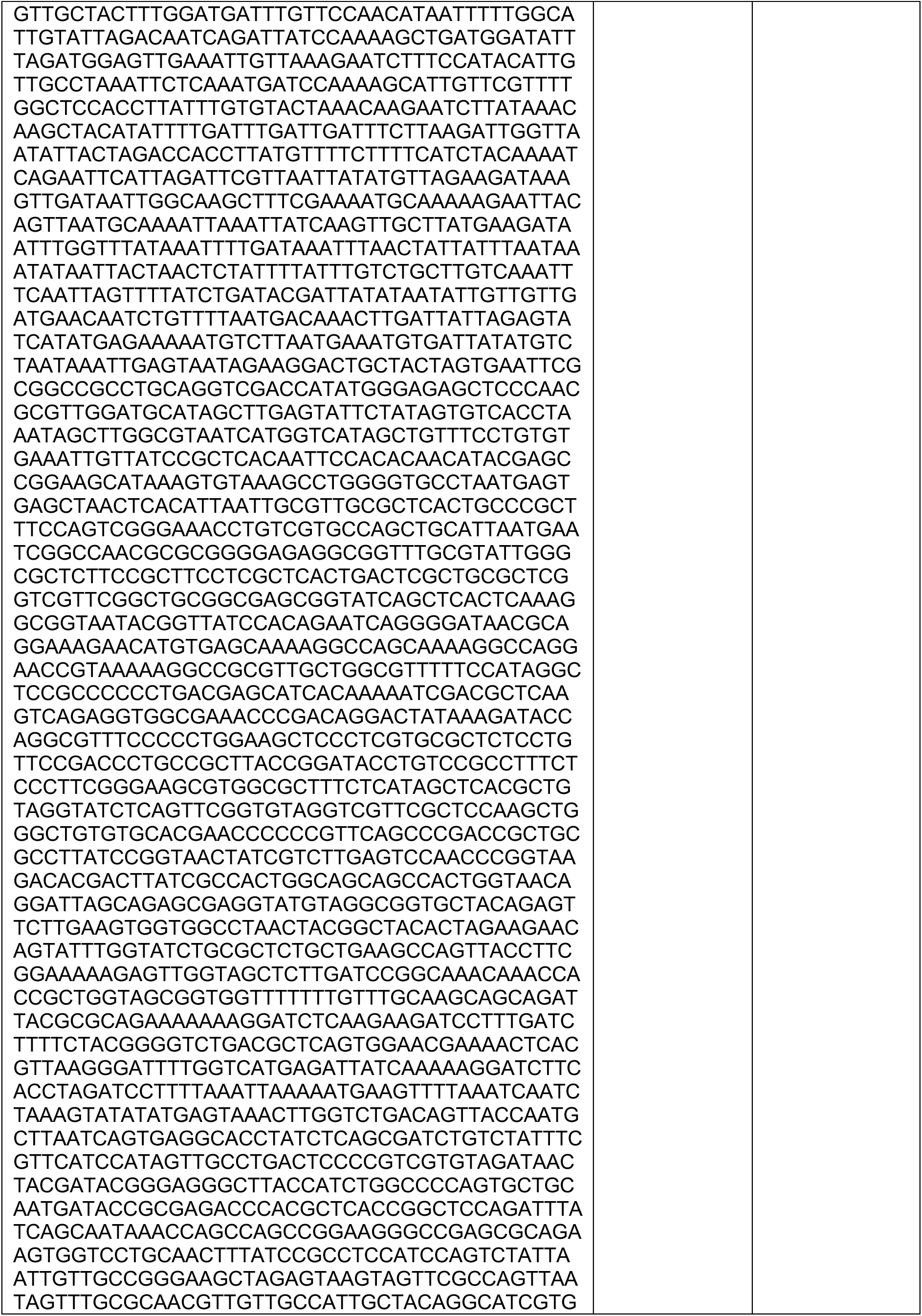
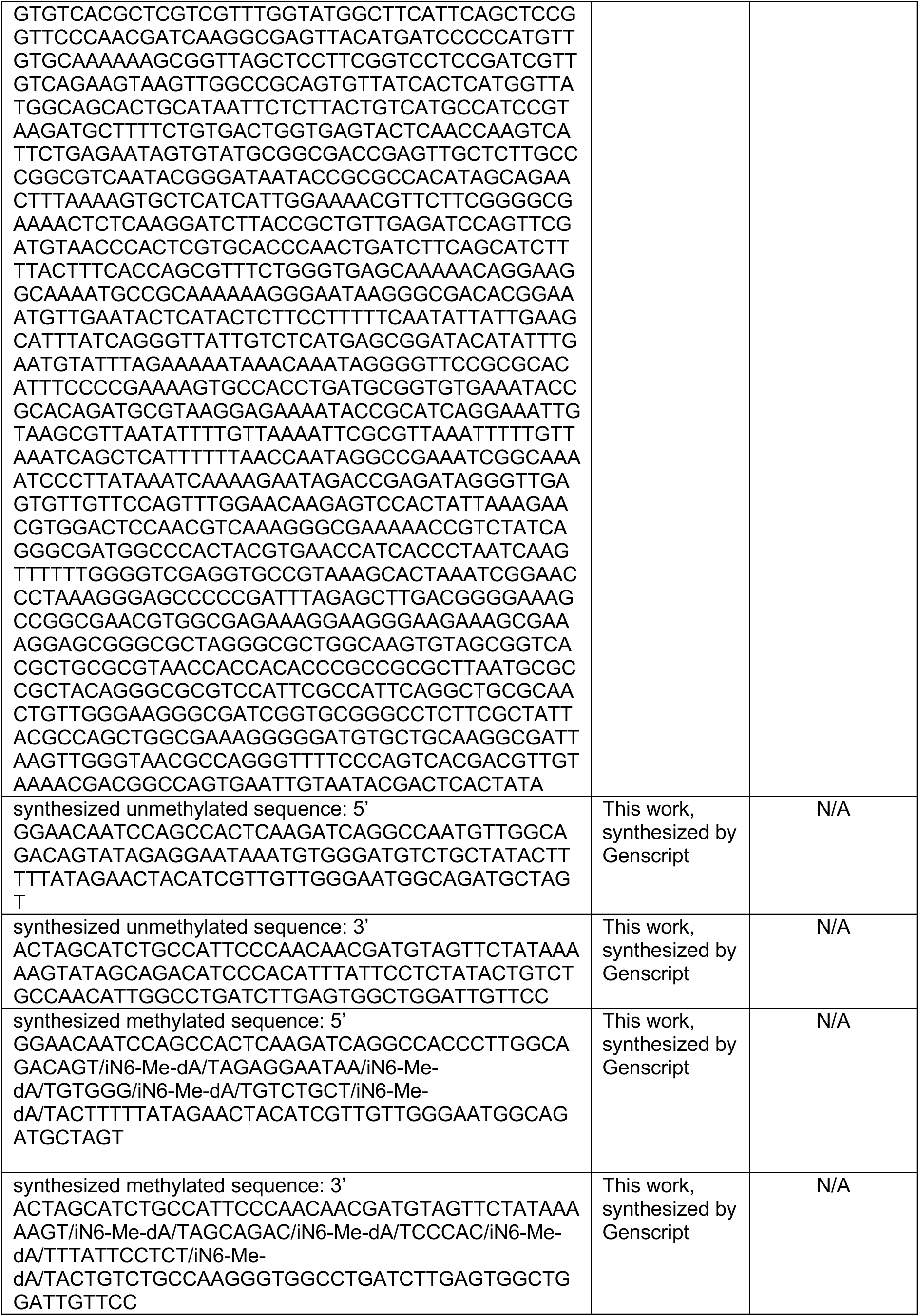
Oligo sequences used for the study. Related to STAR Methods.

## Notes

### Competing Interest Statement

The authors have declared no competing interest.

